# Genomes of Symbiodiniaceae reveal extensive sequence divergence but conserved functions at family and genus levels

**DOI:** 10.1101/800482

**Authors:** Raúl A. González-Pech, Yibi Chen, Timothy G. Stephens, Sarah Shah, Amin R. Mohamed, Rémi Lagorce, Debashish Bhattacharya, Mark A. Ragan, Cheong Xin Chan

**Affiliations:** Institute for Molecular Bioscience, The University of Queensland, Brisbane, QLD 4072, Australia; Commonwealth Scientific and Industrial Research Organisation (CSIRO) Agriculture and Food, Queensland Bioscience Precinct, St Lucia, QLD 4072, Australia; École Polytechnique Universitaire de l’Université de Nice, Université Nice-Sophia-Antipolis, Nice, Provence-Alpes-Côte d’Azur 06410, France; Department of Biochemistry and Microbiology, Rutgers University, New Brunswick, NJ 08901, U.S.A.; School of Chemistry and Molecular Biosciences, The University of Queensland, Brisbane, QLD 4072, Australia

## Abstract

Dinoflagellates of the family Symbiodiniaceae (Order Suessiales) are predominantly symbiotic, and many are known for their association with corals. The genetic and functional diversity among Symbiodiniaceae is well acknowledged, but the genome-wide sequence divergence among these lineages remains little known. Here, we present *de novo* genome assemblies of five isolates from the basal genus *Symbiodinium*, encompassing distinct ecological niches. Incorporating existing data from Symbiodiniaceae and other Suessiales (15 genome datasets in total), we investigated genome features that are common or unique to these Symbiodiniaceae, to genus *Symbiodinium*, and to the individual species *S. microadriaticum* and *S. tridacnidorum*. Our whole-genome comparisons reveal extensive sequence divergence, with no sequence regions common to all 15. Based on similarity of *k*-mers from whole-genome sequences, the distances among *Symbiodinium* isolates are similar to those between isolates of distinct genera. We observed extensive structural rearrangements among symbiodiniacean genomes; those from two distinct *Symbiodinium* species share the most (853) syntenic gene blocks. Functions enriched in genes core to Symbiodiniaceae are also enriched in those core to *Symbiodinium.* Gene functions related to symbiosis and stress response exhibit similar relative abundance in all analysed genomes. Our results suggest that structural rearrangements contribute to genome sequence divergence in Symbiodiniaceae even within a same species, but the gene functions have remained largely conserved in Suessiales. This is the first comprehensive comparison of Symbiodiniaceae based on whole-genome sequence data, including comparisons at the intra-genus and intra-species levels.

## Introduction

Symbiodiniaceae is a family of dinoflagellates (Order Suessiales) that diversified largely as symbiotic lineages, many of which are crucial symbionts for corals. However, the diversity of Symbiodiniaceae extends beyond symbionts of diverse coral reef organisms, to other putative parasitic, opportunistic and free-living forms^1–4^. Genetic divergence among Symbiodiniaceae is known to be extensive, in some cases comparable to that among members of distinct dinoflagellate orders^5^, prompting the recent systematic revision as the family Symbiodiniaceae, with seven delineated genera^6^.

Conventionally, genetic divergence among Symbiodiniaceae has been estimated based on sequence-similarity comparison of a few conserved marker genes. An earlier comparative study using predicted genes from available transcriptome and genome data revealed that functions pertinent to symbiosis are common to all Symbiodiniaceae, but the differences in gene-family number among the major lineages are possibly associated with adaptation to more-specialised ecological niches^7^. A recent investigation^8^ revealed little similarity between the whole-genome sequences of a symbiotic and a free-living *Symbiodinium* species. However, whether this sequence divergence is an isolated case, or is associated with the distinct lifestyles, remains to be investigated using more genome-scale data. In cases such as this, intra-genus and/or intra-species comparative studies may yield novel insights into the biology of Symbiodiniaceae. For instance, a transcriptomic study of four species (with multiple isolates per species) of *Breviolum* (formerly Clade B) revealed differential gene expression that is potentially associated with their prevalence in the host^9^. Comparison of genome data from multiple isolates of the same genus, and/or of the same species, would allow for identification of the molecular mechanisms that underpin the diversification of Symbiodiniaceae at a finer resolution.

In this study, we generated *de novo* genome assemblies from five isolates of *Symbiodinium* (the basal genus of Symbiodiniaceae), encompassing distinct ecological niches (free-living, symbiotic and opportunistic), including two distinct isolates of *Symbiodinium microadriaticum*. Comparing these genomes against those available from other *Symbiodinium*, other Symbiodiniaceae and the outgroup species *Polarella glacialis* (15 datasets in total), we investigated genome features that are common or unique to the distinct lineages within a single species, within a single genus, and within Family Symbiodiniaceae. This is the most comprehensive comparative analysis to date of Symbiodiniaceae based on whole-genome sequence data.

## Results

### Genome sequences of Symbiodiniaceae

We generated draft genome assemblies *de novo* for *Symbiodinium microadriaticum* CassKB8, *Symbiodinium microadriaticum* 04-503SCI.03, *Symbiodinium necroappetens* CCMP2469, *Symbiodinium linucheae* CCMP2456 and *Symbiodinium pilosum* CCMP2461. These five assemblies, generated using only short-read sequence data, are of similar quality to previously published genomes of Symbiodiniaceae (Table 1 and Supplementary Table 1). The number of assembled scaffolds ranges from 37,772 for *S. linucheae* to 104,583 for *S. necroappetens*; the corresponding N50 scaffold lengths are 58,075 and 14,528 bp, respectively. The fraction of the genome recovered in the assemblies ranged from 54.64% (*S. pilosum*) to 76.26% (*S. necroappetens*) of the corresponding genome size estimated based on *k*-mers (Supplementary Table 2). The overall G+C content of all analysed *Symbiodinium* genomes is ~50% (Supplementary Figure 1), with the lowest (48.21%) in *S. pilosum* CCMP2461 and the highest (51.91%) in *S. microadriaticum* CassKB8.

**Table 1.** *Symbiodinium* isolates for which genome data were generated and genome assembly statistics. Details on the *Symbiodinium* isolates for which genome data were generated in this study, and their corresponding genome assembly statistics.

| Isolate details/<br>assembly statistic | *S. microadriaticum* |  | *S. necroappetens* | *S. linuchae* | *S. pilosum* |
| --- | --- | --- | --- | --- | --- |
|  | CassKB8 | 04-503SCI.03 | CCMP2469 | CCMP2456 | CCMP2461 |
| *ITS2*-subtype | A1 | A1 | A13 | A4 | A2 |
| Lifestyle | Symbiotic | Symbiotic | Opportunistic | Symbiotic | Free-living |
| Host | *Cassiopea* sp.<br>(jellyfish) | *Orbicella*<br>*faveolata*<br>(stony coral) | *Condylactis*<br>*gigantea*<br>(anemone) | *Plexaura*<br>*homamalla*<br>(octocoral) | *Zoanthus*<br>*sociatus*<br>(zoanthid) |
| Collection site | Hawaii<br>(Pacific) | Florida<br>(Atlantic) | Jamaica<br>(Caribbean) | Bermuda<br>(Atlantic) | Jamaica<br>(Caribbean) |
| Overall G+C (%) | 51.91 | 50.46 | 50.85 | 50.36 | 48.21 |
| Number of scaffolds | 67,937 | 57,558 | 104,583 | 37,772 | 48,302 |
| Assembly length (bp) | 813,744,491 | 775,008,844 | 767,953,253 | 694,902,460 | 1,089,424,773 |
| N50 scaffold length<br>(bp) | 42,989 | 49,975 | 14,528 | 58,075 | 62,444 |
| Max. scaffold length<br>(Mbp) | 0.38 | 1.08 | 1.34 | 0.46 | 1.34 |
| Number of contigs | 167,159 | 162,765 | 157,685 | 141,380 | 142,969 |
| N50 contig length (bp) | 10,400 | 11,136 | 11,420 | 11,147 | 17,506 |
| Max. contig length<br>(Mbp) | 0.15 | 1.05 | 1.34 | 0.19 | 1.34 |
| Gap (%) | 1.15 | 1.44 | 0.56 | 1.35 | 0.79 |
| Estimated genome size<br>(bp) | 1,120,150,369 | 1,052,668,212 | 1,007,022,374 | 914,781,885 | 1,993,912,458 |
| Assembled fraction of<br>genome (%) | 72.65 | 73.62 | 76.26 | 75.96 | 54.64 |

For a comprehensive comparison, we included in our analysis all available genome data from Symbiodiniaceae and the outgroup species of *Polarella glacialis* (Supplementary Table 1). These data comprise nine *Symbiodinium* isolates (three of the species *S. microadriaticum* and two of *S. tridacnidorum*), *Breviolum minutum*, two *Cladocopium* isolates, *Fugacium kawagutii*, and two *Polarella glacialis* isolates^8,10–14^ (*i.e.* a total of 15 datasets of Suessiales, of which 13 are of Symbiodiniaceae); we used the revised genome assemblies from Chen *et al.*^15^ where applicable. Of the 15 genome assemblies, four were generated using both short- and long-read data (those of *S. natans* CCMP2548, *S. tridacnidorum* CCMP2592 and the two *P. glacialis* isolates)^8,14^; all others were generated largely using short-read data.

### Isolates of Symbiodiniaceae and *Symbiodinium* exhibit extensive genome divergence

We assessed genome-sequence similarity based on pairwise whole-genome sequence alignment (WGA). In each pairwise comparison, we assessed the overall percentage of the query genome sequence that aligned to the reference (*Q*), and the average percent identity of the reciprocal best one-to-one aligned sequences (*I*); see Methods for detail. Our results revealed extensive sequence divergence among the compared genomes at the order (Suessiales), family (Symbiodiniaceae) and genus (*Symbiodinium*) levels (Fig. 1A). As expected, the genome-pairs that exhibit the highest sequence similarity are isolates from the same species, *e.g.* between *S. microadriaticum* CassKB8 and 04-503SCI.03 (*Q* = 87.44%, *I* = 99.72%; CassKB8 as query), and between the two *P. glacialis* isolates (*Q* = 97.10%, *I* = 98.59%; CCMP1383 as query). In contrast, genome sequences of the two *S. tridacnidorum* isolates appear more divergent (*Q* = 30.07%, *I* = 87.18%; CCMP2592 as query). Remarkably, some genomes within *Symbiodinium* are as divergent as those of distinct genera: for instance, *Q* = 1.10% and *I* = 91.88% for *S. pilosum* compared against *S. natans* as reference, and *Q* = 1.03% and *I* = 92.15% for *S. tridacnidorum* CCMP2592 against *Cladocopium* sp. C92. The genome sequences of *S. microadriaticum* CCMP2467 share the most genome regions with all analysed isolates (Fig. 1A). When compared against these sequences as reference, we did not recover any genome regions that are conserved (alignment length ≥24 bp, with >70% identity) in all analysed isolates (Fig. 1B). At most, six isolates have genome regions aligned against the reference, all of which belong to the same genus: *S. microadriaticum* CassKB8, *S. microadriaticum* 04-503SCI.03, *S. linucheae*, *S. tridacnidorum* CCMP2592, *S. natans* and *S. pilosum*. However, the total length of the region common in these genomes is only 89 bp (Fig. 1B).

**Fig. 1.**
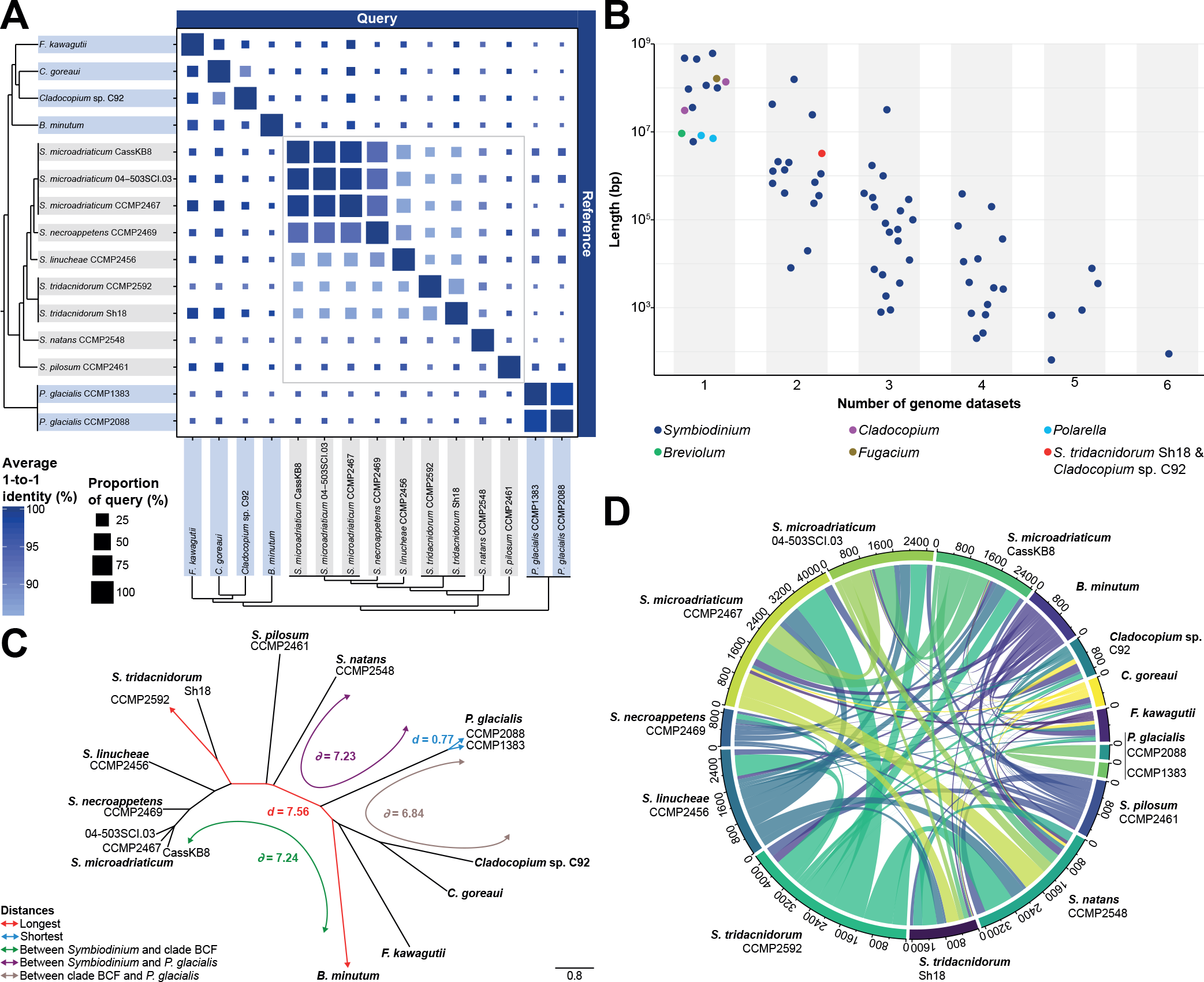
Genome divergence among Symbiodiniaceae. **(A)** Similarity between Symbiodiniaceae (and the outgroup *P. glacialis*) based on pairwise whole-genome sequence alignments. The colour of the square depicts the average percent identity of the best reciprocal one-to-one aligned regions (*I*) between each genome pair and the size of the square is proportional to the percent of the query genome that aligned to the reference (*Q*), as shown in the legend. The tree topologies on the left and bottom indicate the known phylogenetic relationship^6^ among the isolates. Isolates in *Symbiodinium* are highlighted in grey. **(B)** Total sequence length (*y*-axis) of genomic regions aligning to the reference genome assembly of *S. microadriaticum* CCMP2467 shared by different numbers of the datasets used in this study (*x*-axis). Data points represent distinct combinations of datasets, ranging from one (an individual genome dataset) to six (six datasets aligning to the same regions of the reference), and are coloured to show the genera to which they correspond; only one combination includes distinct genera (*S. tridacnidorum* Sh18 and *Cladocopium* sp. C92). **(C)** NJ tree based on 21-mers shared by genomes of Suessiales; branch lengths are proportional to the estimated distances (see Methods). The shortest and longest distances (*d*) in the tree, as well as average distances 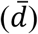 among representative clades are shown following the bottom-left colour code. ‘Clade BCF’: clade including *B. minutum*, *F. kawagutii* and the two *Cladocopium* isolates. **(D)** Number of collinear syntenic gene blocks shared by pairs of genomes of Suessiales. Gene blocks shared by more than two isolates are not shown.

For each possible genome-pair, we also assessed the extent of shared *k*-mers (short, sub-sequences of defined length *k*) between them (optimised *k* = 21; see Methods) from which a pairwise distance (*d*) was derived (Supplementary Table 3). These distances were used to infer the phylogenetic relationship of these genomes as a neighbour-joining (NJ) tree (Fig. 1C) and as a similarity network (Supplementary Figure 2). As shown in Fig. 1C, the most distant genome-pair (*i.e.* the pair with the highest *d*) is *S. tridacnidorum* CCMP2592 and *B. minutum* (*d* = 7.56). *Symbiodinium* isolates are about as distant from the other Symbiodiniaceae 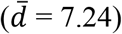 as they are from the outgroup *P. glacialis* 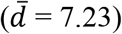. This is surprising, in particular because *P. glacialis* isolates have shorter distances with the other Symbiodiniaceae 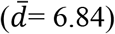 and *Symbiodinium* is considered to be more ancestral than all other genera in Symbiodiniaceae^6^. However, this observation may be biased by the greater representation of *Symbiodinium* isolates compared to any other genera of Symbiodiniaceae. The largest distance among genome-pairs within *Symbiodinium* is between two free-living species, *S. natans* and *S. pilosum* (*d* = 5.64). These two isolates are also the most divergent from all others in the genus (*d* > 4.50 between either of them and any other *Symbiodinium*; Supplementary Table 3). The distance between *S. natans* and *S. pilosum* is similar to that observed between *F. kawagutii* and *C. goreaui* (*d* = 5.74), members of distinct genera. Similar to our WGA results, the shortest distances are between isolates of the same species, *e.g. d* = 0.77 between *P. glacialis* CCMP1383 and CCMP2088, and 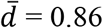 among *S. microadriaticum* isolates. However, the distance between the two *S. tridacnidorum* isolates (CCMP2592 and Sh18; *d* = 2.87) is larger than that between *S. necroappetens* and *S. linucheae* (*d* = 2.66). The divergence among *Symbiodinium* isolates is further supported by the mapping rate of paired reads (Supplementary Figure 3).

We used the same gene-prediction workflow, customised for dinoflagellates, for the five *Symbiodinium* genome studies generated in this study as for the other ten assemblies included in our analyses^14,15^ (Table 1). The number of predicted genes in these genomes ranged between 23,437 (in *S. pilosum* CCMP2461) and 42,652 (in *S. microadriaticum* CassKB8), which is similar to the number of genes (between 25,808 and 45,474) predicted in the other Symbiodiniaceae genomes (Supplementary Table 4). To further assess genome divergence, we identified conserved synteny based on collinear syntenic gene blocks (see Methods). Fig. 1D illustrates the gene blocks shared between any possible genome-pairs; those blocks shared by more than two genomes are not shown. *S. microadriaticum* CCMP2467 and *S. tridacnidorum* CCMP2592 share the most gene blocks (853 implicating 8589 genes). Although the two *P. glacialis* genomes share 346 gene blocks (2524 genes), no blocks were recovered between the genome of either *P. glacialis* isolate and any of *S. microadriaticum* CassKB8, *S. microadriaticum* 04-503SCI.03, *S. necroappetens*, *C. goreaui*, *Cladocopium* sp. C92 or *F. kawagutii*. The collinear gene blocks shared by *P. glacialis* CCMP1383 and *S. microadriaticum* CCMP2467 (3 blocks, 19 genes) represent the most abundant between any *P. glacialis* and any Symbiodiniaceae isolate. Genomes of *S. tridacnidorum* CCMP2592 and *S. natans* more gene blocks (749, with 7290 genes) than any other pair of genomes within Symbiodiniaceae. Although we cannot dismiss the impact of contiguity and completeness of the genome assemblies (Supplementary Table 1, Supplementary Figure 4) on our observations here (and results from the WGA and *k*-mer analyses above), these results provide the first comprehensive overview of genome divergence at the resolution of species, genus and family levels.

### Remnants of transposable elements were lost in more-recently diverged lineages of Symbiodiniaceae

Fig. 2A shows the composition of repeats for each of the 15 genomes. The repeat composition of *P. glacialis* is distinct from that of Symbiodiniaceae genomes, largely due to the known prevalence of simple repeats^8,14^. Long interspersed nuclear elements (LINEs) in Symbiodiniaceae and in *P. glacialis* are highly diverged, with Kimura distance centred between 15 and 40; these elements likely represent remnants of LINEs from an ancient burst pre-dating the diversification of Suessiales^8,11,14^. Interestingly, the proportion of these elements is substantially larger in the genomes of *Symbiodinium* (the basal lineage) and *P. glacialis* (the outgroup) than in those of other Symbiodiniaceae (Fig. 2B). For instance, LINEs comprise between 74.10 Mbp (*S. tridacnidorum* Sh18) and 96.9 Mbp (*S. linucheae*) in each of the *Symbiodinium* genomes, except for those in *S. pilosum* that cover almost twice as much (171.31 Mbp). In comparison, LINEs cover on average 7.49 Mbp in the genomes of other Symbiodiniaceae (Supplementary Figure 5**Error! Reference source not found.**). This result suggests that the remnants of LINEs were lost in the more-recently diverged lineages of Symbiodiniaceae.

**Fig. 2.**
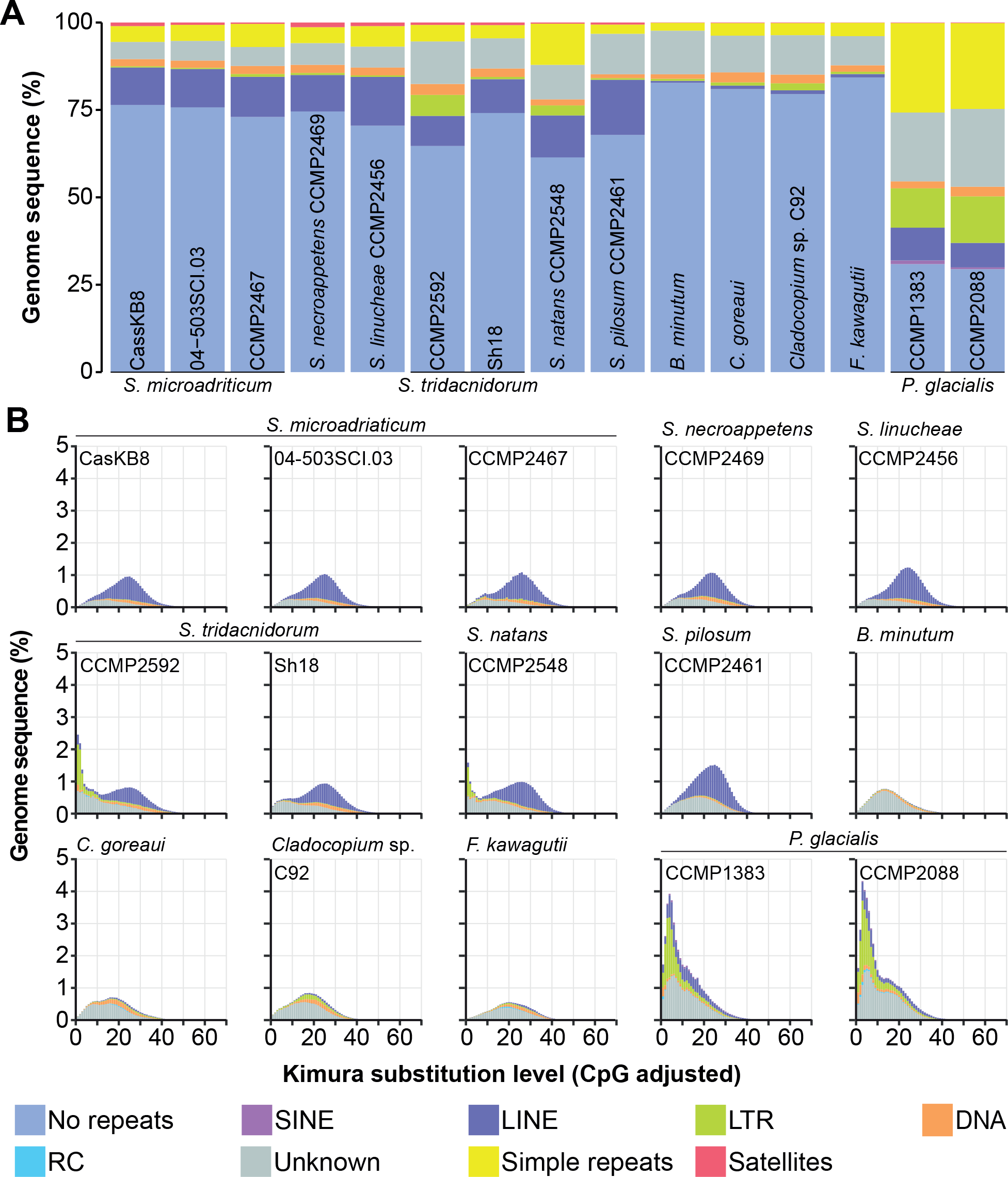
Repeat composition of Suessiales genomes. **(A)** Percentage of sequence regions comprising the major classes of repetitive elements, shown for each genome assembly analysed in this study. **(B)** Interspersed repeat landscape for each assembled genome. Both **(A)** and **(B)** follow the colour code shown in the bottom legend.

The genome of the free-living *S. pilosum* presents an outlier among the *Symbiodinium* genomes. In addition to the nearly two-fold increased abundance of LINEs, the estimated genome size for *S. pilosum* (1.99 Gbp) is also nearly two-fold larger than the estimate for any other *Symbiodinium* genome (Supplementary Table 2). This suggests whole-genome duplication or potentially a more-dominant diploid stage, but we found no evidence to support either scenario (Supplementary Figure 6). The prevalence of repetitive regions in *S. pilosum*, however, would explain in part why the total assembled bases of the genome constitute only 54.64% of the estimated genome size (Supplementary Table 1).

### Diversity of gene features within Suessiales

Differences among predicted genes of Symbiodiniaceae have been attributed to phylogenetic relationship and to the implementation of distinct gene prediction approaches^15^. Our Principal Component Analysis (PCA), based on metrics of consistently predicted genes (Supplementary Table 4), revealed substantial variation within the genus *Symbiodinium* (Fig. 3). We noticed that the observed variation can be associated with three main factors: (1) phylogenetic relationship, (2) the type of sequence data used for genome assembly and the consequent assembly quality, and (3) lifestyle of the isolates. The variation resulting from the phylogenetic relationship among the genomes is illustrated by the separation of the distinct genera along PC2 (explaining 24.82% of the variance). The metrics contributing the most to PC2 are associated with proportion of splice donors and acceptors (Supplementary Figure 7). The type of sequence data used for genome assembly and assembly quality are reflected along PC1 (explaining 42.79% of the variance). For instance, taxa for which hybrid assemblies were made (those incorporating both short-read and long-read sequence data), *i.e.* the free-living *S. natans* and *P. glacialis*, and the symbiotic *S. tridacnidorum* CCMP2592, are distributed between −4.5 and 0.1 along PC1. The distribution of the symbiotic *Symbiodinium* is limited (between 0.5 and 1.5 of PC1), with the exception of the two *S. tridacnidorum* isolates, for which the genome assemblies are of distinct quality (*i.e.* the high-quality hybrid assembly of CCMP2592 compared to the draft assembly of Sh18 that is fragmented and incomplete; Supplementary Table 1 and Supplementary Figure 4). In addition, the opportunistic *S. necroappetens* and free-living *S. pilosum* are distributed at >2 along PC1. These observations suggest that the distinct lifestyles may contribute to differences in gene architecture.

**Fig. 3.**
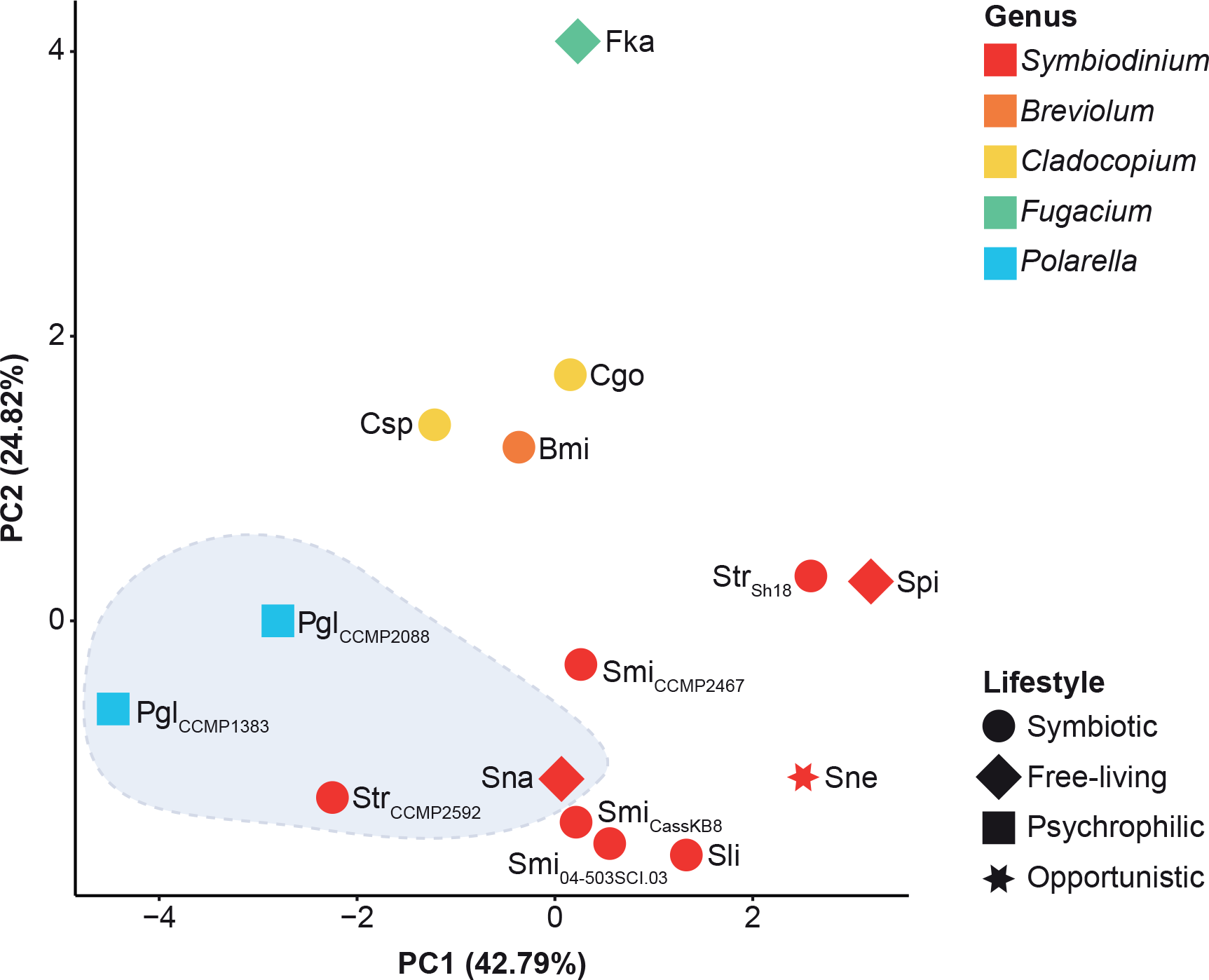
PCA of gene features in Symbiodiniaceae. PCA displaying the variation of predicted genes among the analysed genomes based on gene metrics (Supplementary Table 4). Data points are coloured by genus and shaped by lifestyles according to the legends to the right. Data points enclosed in a light blue area correspond to isolates with hybrid genome assemblies. Smi: *S. microadriaticum*, Sne: *S. necroappetens*, Sli: *S. linucheae*, Str: *S. tridacnidorum*, Sna: *S. natans*, Spi: *S. pilosum*, Bmi: *B. minutum*, Cgo: *C. goreaui*, Csp: *Cladocopium* sp. C92, Fka: *F. kawagutii*, Pgl: *P. glacialis*. Isolate name is shown in subscript for those species with more than one isolate.

The predicted coding sequences (CDS) among *Symbiodinium* taxa exhibit biases in nucleotide composition of codon positions (Supplementary Figure 8) and in codon usage (Supplementary Figure 9). The G+C content among CDS (Supplementary Table 4) and among third codon positions (Supplementary Figure 8) varies slightly, but is generally higher relative to the overall G+C content (Supplementary Figure 1, Supplementary Table 1). This is consistent with the results previously reported for genomes and transcriptomes of Symbiodiniaceae^7,16^. Of all *Symbiodinium* isolates, *S. microadriaticum* CassKB8 and 04-503SCI.03 have the most CDS with a strong codon preference; *S. microadriaticum* CCMP2467 has the least (Supplementary Figure 9). These observations highlight the genetic variation within a single genus, and within a single species.

### Gene families of Symbiodiniaceae

Using all 555,682 predicted protein sequences from the 15 genomes, we inferred 42,539 homologous sets (of size ≥ 2; see Methods); here we refer to these sets as gene families. Of the 42,539 families, 18,453 (43.38%) contain genes specific to Symbiodiniaceae (Fig. 4). Interestingly, more (8828) gene families are specific to sequenced isolates of *Symbiodinium* than to sequenced isolates of the other Symbiodiniaceae combined (2043 specific to *Breviolum*, *Cladocopium* and *Fugacium* isolates). Although the simplest explanation is that substantially more gene families have been gained (or preserved) in *Symbiodinium* than in the other three genera, we cannot dismiss potential biases caused by our more-comprehensive taxon sampling for this genus. In contrast, a previous study reported substantially more gene families specific to the clade encompassing *Breviolum*, *Cladocopium* and *Fugacium* (26,474) than specific to *Symbiodinium* (3577)^7^. It is difficult to compare these two results because the previous study used predominantly transcriptomic data (which are fragmented and include transcript isoforms), proteins predicted with distinct and inconsistent methods, and a different approach to delineate gene families.

**Fig. 4.**
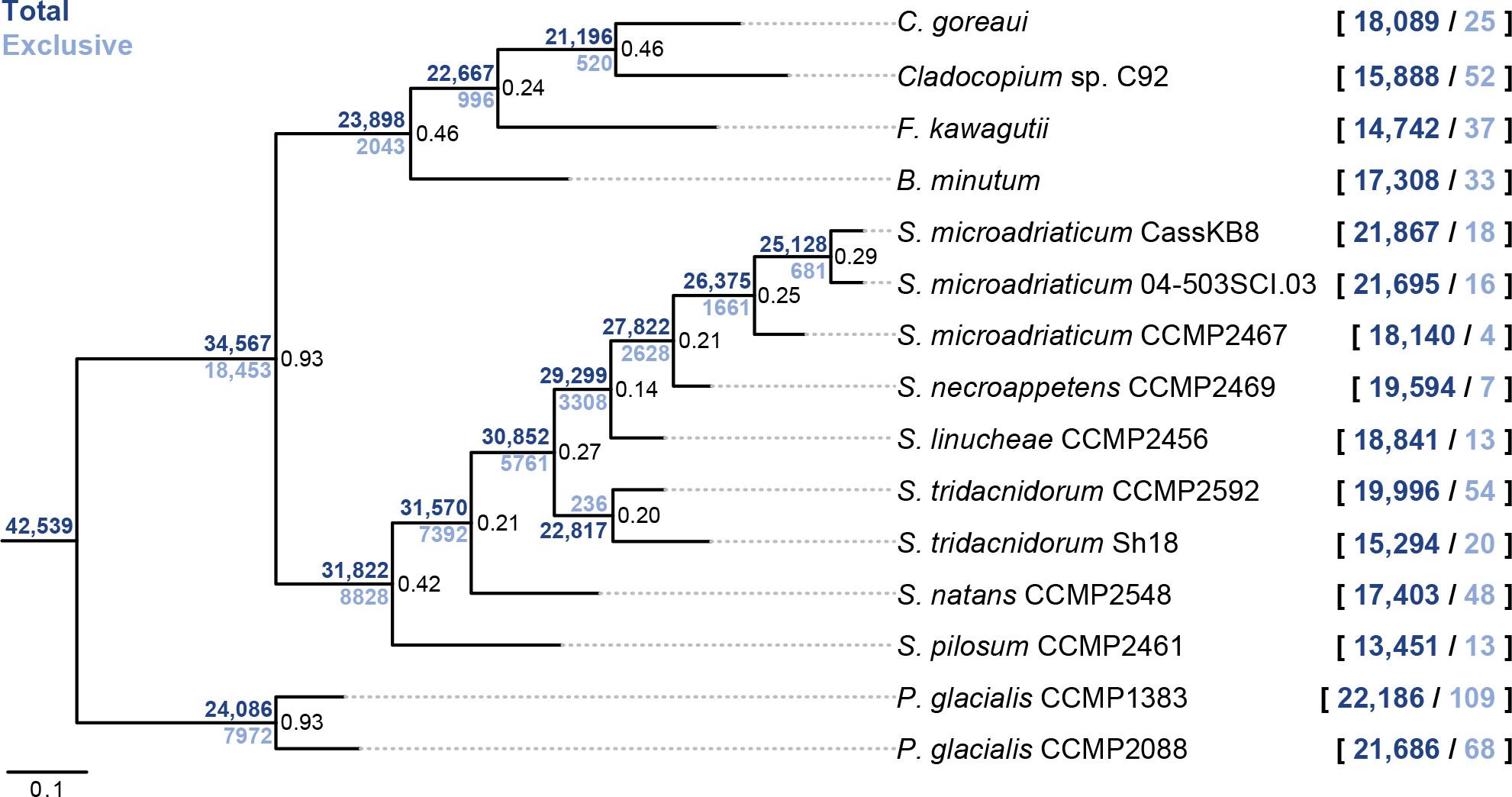
Number of gene families along the phylogeny of Symbiodiniaceae. Species tree inferred based on 28,116 gene families containing at least 4 genes from any Suessiales isolate using STAG^67^ and STRIDE^68^ (part of the conventional OrthoFinder pipeline^66^), rooted with *P. glacialis* as outgroup. At each node, the total number of families that include genes from one or more diverging isolates is shown in dark blue, those exclusive to one or more diverging isolates in light blue. The numbers shown for each isolate (on the right) represent numbers of gene families that include genes from (dark blue) and exclusive to (light blue) that isolate. The proportion of gene trees supporting each node is shown. Branch lengths are proportional to the number of substitutions per site.

Of all families, 2500 (5.88%) contain genes from all 15 Suessiales isolates; 4677 (10.9%) represent 14 or more isolates. We consider these 4677 as the core gene families to Suessiales. Only 406 gene families are exclusive and common to all 13 Symbiodiniaceae isolates; 914 represent 12 or more isolates. Similarly, 193 are exclusive and common to all nine *Symbiodinium* isolates; 539 represent eight or more isolates. We define these 914 and 539 families as the core gene families for Symbiodiniaceae and for *Symbiodinium*, respectively.

Despite the variable quality and completeness of the genome assemblies analysed here (Supplementary Table 1, Supplementary Figure 4), we consider these results more reliable than those based largely on transcriptome data^7^, in which transcript isoforms, in addition to quality and completeness of the datasets, can result in overestimation of gene numbers and introduce noise and bias to the data. The smaller number of gene families shared among Symbiodiniaceae found here (*i.e.* 18,453 compared to 76,087 in the earlier study^7^) likely reflects our more-conservative approach based on whole-genome sequenced data. Nonetheless, our observations support the notion that evolution of gene families has contributed to the diversification of Symbiodiniaceae^7^.

### Core genes of Symbiodiniaceae and of *Symbiodinium* encode similar functions

To identify gene functions characteristic of Symbiodiniaceae and *Symbiodinium*, we carried out enrichment analyses based on Gene Ontology (GO)^17^ of the annotated gene functions in the corresponding core families. Among the core genes of Symbiodiniaceae, the most significantly overrepresented GO terms relate to retrotransposition, components of the membrane (including ABC transporters), cellulose binding, and reduction and oxidation reactions of the electron transport chain (Supplementary Table 5). Retrotransposition has been shown to contribute to gene-family expansion and changes in the gene structure of Symbiodiniaceae^8,18^. The enrichment of this function in Symbiodiniaceae may be due to a common origin of genes that encode remnant protein domains from past retrotransposition events (*e.g.* genes encoding reverse transcriptase, as previously reported^8^). Proteins integrated in the cell membrane are relevant to symbiosis^19,20^. For instance, ABC transporters may play a major role in the exchange of nutrients between host and symbiotic Symbiodiniaceae^21^. The enrichment of cellulose-binding function may be related to the changes in the cell wall during the transition between the mastigote and coccoid stages common in symbiotic Symbiodiniaceae^22^. The overrepresentation of electron transport chain functions may be associated with the acclimation of Symbiodiniaceae to different light conditions and/or to adjustments of the thylakoid membrane composition to prevent photoinhibition under stress^23,24^.

Similarly, among core genes of *Symbiodinium*, the most significantly enriched functions are related to retrotransposition (Supplementary Table 6). This is likely a reflection of the higher content of LINEs in *Symbiodinium* genomes (and perhaps also of LTRs in *S. tridacnidorum* CCMP2592 and *S. natans* CCMP2548) compared to the other Symbiodiniaceae isolates (Fig. 2 and Supplementary Figure 5). Nevertheless, the presence of retrotransposition among the functions overrepresented in the cores of both Symbiodiniaceae and *Symbiodinium* supports the notion of substantial divergence, potentially result of pseudogenisation or neofunctionalisation, accumulated between gene homologs that prevents the clustering of these homologs within the same gene family^7,8^.

### Functions related to symbiosis and stress response are conserved in Suessiales

We further examined the functions annotated for the predicted genes of all 15 Suessiales isolates based on the annotated GO terms and protein domains. A recent study, focusing on the transcriptomic changes in *Cladocopium* sp. following establishment of symbiosis with coral larvae^21^, complied a list of symbiosis-related gene functions in Symbiodiniaceae. We searched for these functions, and found that they are conserved in Symbiodiniaceae regardless of the lifestyle (*e.g.* the free-living *S. natans*, *S. pilosum* and *F. kawagutii*, or the opportunistic *S. necroappetens*), and even in the outgroup *P. glacialis* (Fig. 5). This result supports the notion that genomes of dinoflagellates encode gene functions conducive to adaptation to a symbiotic lifestyle^10^. However, we observed a trend of reduced abundance of these functions in genes of *B. minutum*, *C. goreaui* and *Cladocopium* sp. C92, with the exception of genes encoding ankyrin and tetratricopeptide repeat domains. Although multiple Pfam domains of ankyrin or tetratricopeptide repeats exist, all isolates exhibit consistently higher abundance for specific types (PF12796 and PF13424, respectively). Interestingly, despite the presence of ABC transporters in the enriched functions of the core genes of Symbiodiniaceae (Supplementary Table 5), they appear to occur in low abundance.

**Fig. 5.**
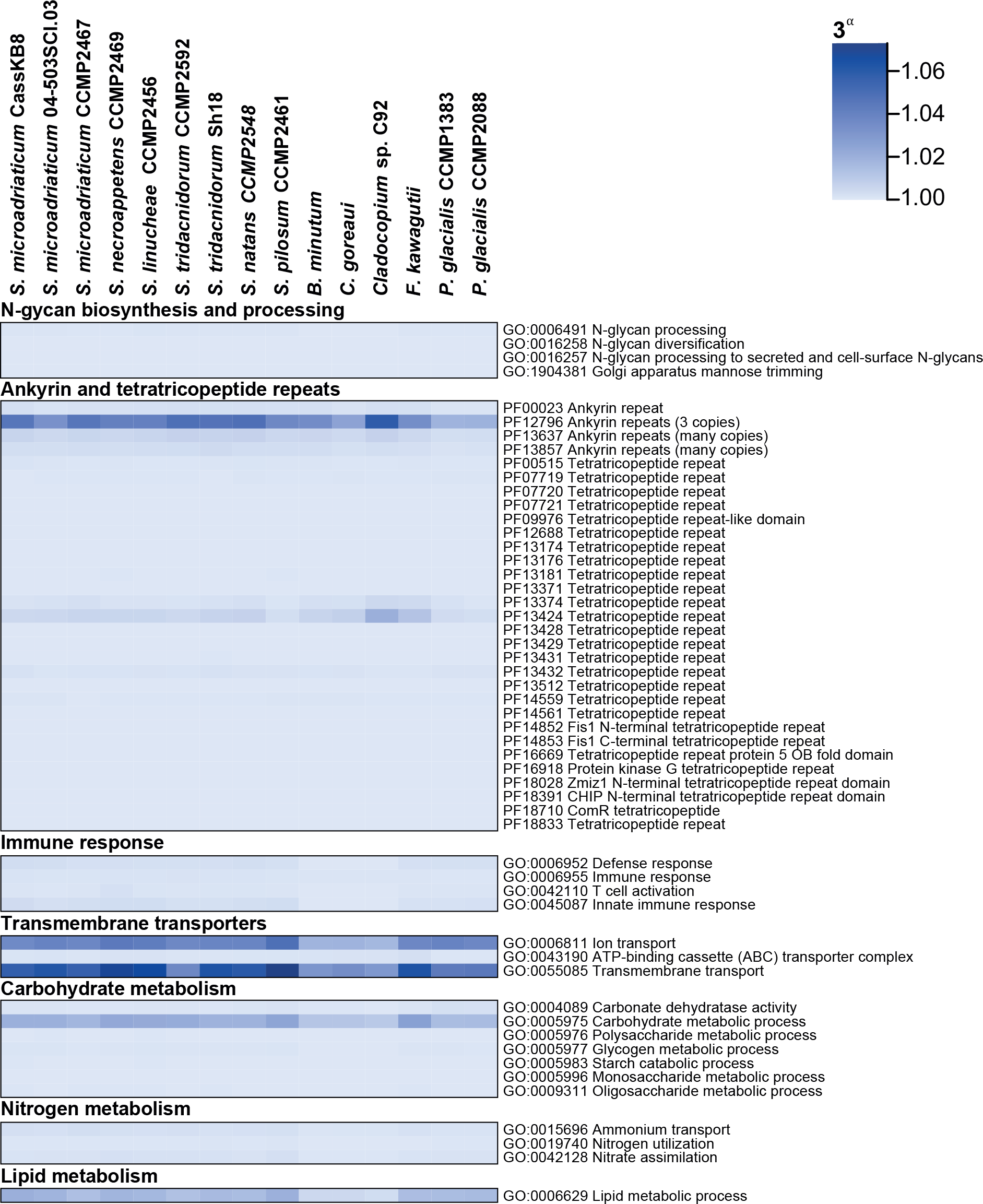
Relative abundance of symbiosis-related functions in genes of Suessiales. Heat map showing the relative abundance (α) of GO terms (relative to the total number of genes) and protein domains (relative to the total number of identified domains) that are related to symbiosis shown for each genome. The transformed values of α are shown in the form of 3^α^.

The abundance of functions associated with response to distinct types of stress, cell division, DNA damage repair, photobiology and motility also appear to be conserved across Suessiales (Fig. 6). The abundance of genes annotated with DNA repair functions is consistent with the previously reported overrepresentation of these functions in genomes and transcriptomes of Suessiales^7^ and the presence of gene orthologs involved in a wide range of DNA damage responses in dinoflagellates^25^. Likewise, the relatively high abundance of functions related to DNA recombination may represent further support for the potential of sexual reproduction in these dinoflagellates^11,26^, and for the contribution of sexual recombination to genetic diversity of Symbiodiniaceae^27–31^. Moreover, the higher abundance of a cold-shock DNA-binding domain and bacteriorhodopsin in *P. glacialis* compared to the Symbiodiniaceae isolates highlights the adaptation of this species to extreme cold and low-light environments, and is consistent with the highly duplicated genes encoding these functions in *P. glacialis* genomes^14^.

**Fig. 6.**
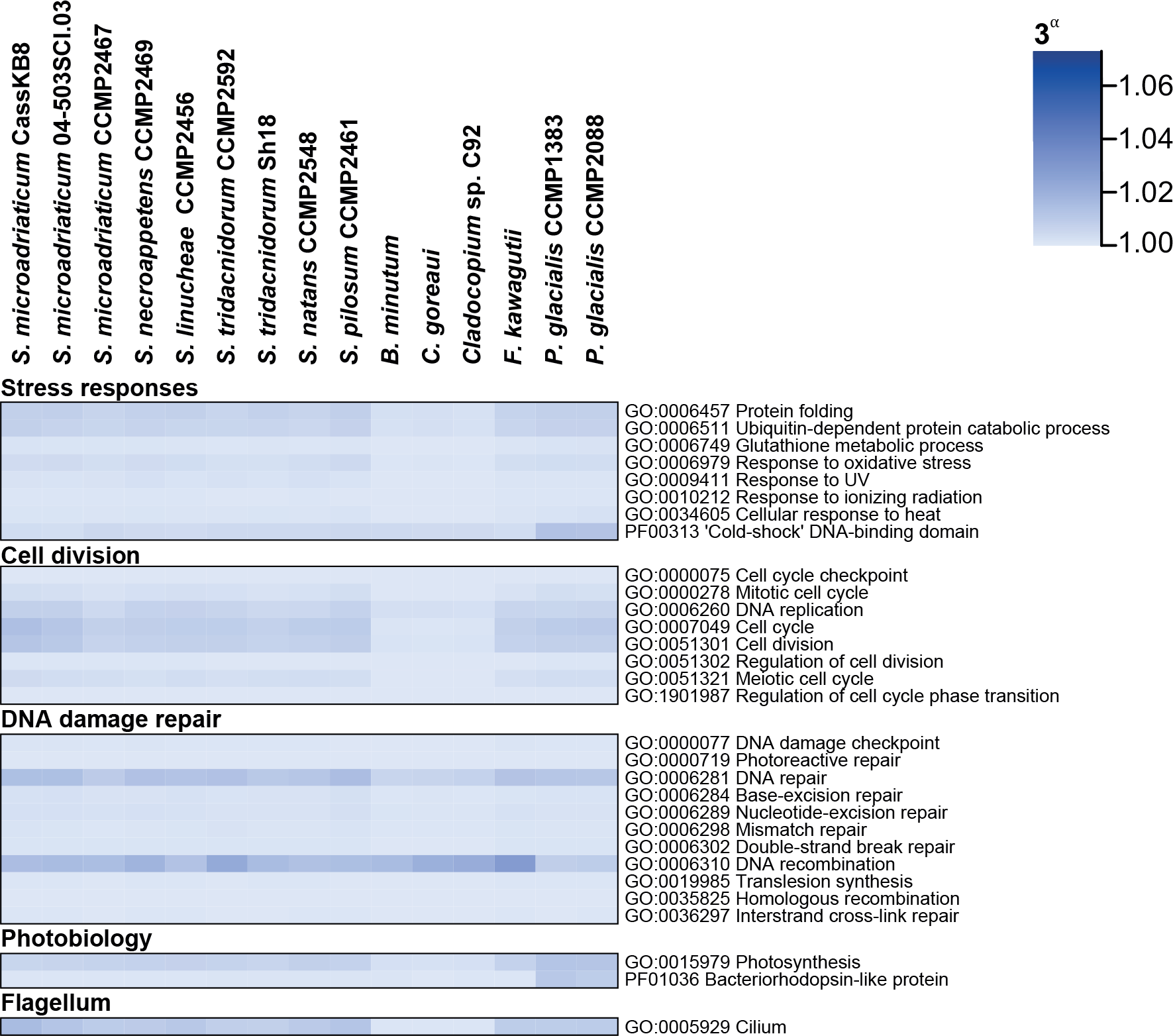
Relative abundance of selected functions in genes of Suessiales. Heat map showing the relative abundance (α) of GO terms (relative to the total number of genes) and protein domains (relative to the total number of identified domains) that are associated with key functions shown for each genome. The transformed values of α are shown in the form of 3^α^.

## Discussion

Our results suggest that whereas gene functions appear to be largely conserved across isolates from the same order (Suessiales), family (Symbiodiniaceae) and genus (*Symbiodinium*), there is substantial genome-sequence divergence among these isolates. However, what drives this divergence remains an open question. Although sexual recombination probably contributes to the extensive genetic diversity in Symbiodiniaceae^27–31^, its limitation to homologous regions renders its contribution as the sole driver of divergence unlikely. The evolutionary transition from a free-living to a symbiotic lifestyle can contribute to the loss of conserved synteny as consequence of large- and small-scale structural rearrangements^16,32,33^. The enhanced activity of mobile elements in the early stages of this transition can further disrupt synteny, impact gene structure and accelerate mutation rate^34,35^. However *S. natans* and *S. pilosum*, for which the free-living lifestyle has been postulated to be ancestral^8^, are still quite divergent from each other (Fig. 1). Ancient events, such as geological changes or emergence of hosts, are thought to influence diversification of Symbiodiniaceae^6,36,37^ and may help explain the divergence of the extant lineages. For example, in a hypothetical scenario, drastic changes in environmental conditions could have split the ancestral Symbiodiniaceae population into multiple sub-populations with very small population sizes. This would have enabled rapid divergence among the sub-populations that, in turn, could have evolved and diversified independently into the extant taxa.

Although genome data generally provide a comprehensive view of gene functions, we cannot dismiss artefacts that may have been introduced by the type of sequence data used to generate the genome assemblies analysed here. Genes encoding functions critical to dinoflagellates often occur in multiple copies, and those of Symbiodiniaceae are no exceptions^8,10,14^. Incorporation of long-read sequence data in the genome assembly is important to resolve repetitive elements (including genes occurring in multiple copies) and allow for more-accurate analysis of abundance or enrichment of gene functions. On the other hand, accurate inference of gene families can be challenging especially for gene homologs with an intricate evolutionary history. Moreover, a good taxa representation can aid the inference of homology^38,39^. Data that better resolve multi-copy genes (*e.g.* through the incorporation of long-read sequences in the assembly process^8^) will allow better understanding of gene loss and innovation along the genome evolution of Symbiodiniaceae.

This work reports the first whole-genome comparison at multiple taxonomic levels within dinoflagellates: within Order Suessiales, within Family Symbiodiniaceae, within Genus *Symbiodinium,* and separately for the species *S. microadriaticum* and *S. tridacnidorum*. We show that whereas genome sequences can diverge substantially among Symbiodiniaceae, gene functions nonetheless remain largely conserved even across Suessiales. Our understanding of the evolution of this remarkably divergent family would benefit from more-narrowly scoped studies at the intra-generic and intra-specific levels. Even so, our work demonstrates the value of comprehensive surveys to unveil macro-evolutionary processes that led to the diversification of Symbiodiniaceae.

## Methods

### *Symbiodinium* cultures

Single-cell monoclonal cultures of *S. microadriaticum* CassKB8 and *S. microadriaticum* 04-503SCI.03 were acquired from Mary Alice Coffroth (Buffalo University, New York, USA), and those of *S. necroappetens* CCMP2469, *S. linucheae* CCMP2456 and *S. pilosum* CCMP2461 were purchased from the National Center for Marine Algae and Microbiota at the Bigelow Laboratory for Ocean Sciences, Maine, USA (Table 1). The cultures were maintained in multiple 100-mL batches (in 250-mL Erlenmeyer flasks) in f/2 (without silica) medium (0.2 mm filter-sterilized) under a 14:10 h light-dark cycle (90 μE/m^2^/s) at 25 ºC. The medium was supplemented with antibiotics (ampicillin [10 mg/mL], kanamycin [5 mg/mL] and streptomycin [10 mg/mL]) to reduce bacterial growth.

### Nucleic acid extraction

Genomic DNA was extracted following the 2×CTAB protocol with modifications. *Symbiodinium* cells were first harvested during exponential growth phase (before reaching 106 cells/mL) by centrifugation (3000 *g*, 15 min, room temperature (RT)). Upon removal of residual medium, the cells were snap-frozen in liquid nitrogen prior to DNA extraction, or stored at −80 °C. For DNA extraction, the cells were suspended in a lysis extraction buffer (400 μL; 100 mM Tris-Cl pH 8, 20 mM EDTA pH 8, 1.4 M NaCl), before silica beads were added. In a freeze-thaw cycle, the mixture was vortexed at high speed (2 min), and immediately snap-frozen in liquid nitrogen; the cycle was repeated 5 times. The final volume of the mixture was made up to 2% w/v CTAB (from 10% w/v CTAB stock; kept at 37 °C). The mixture was treated with RNAse A (Invitrogen; final concentration 20 μg/mL) at 37 °C (30 min), and Proteinase K (final concentration 120 μg/mL) at 65 °C (2 h). The lysate was then subjected to standard extractions using equal volumes of phenol:chloroform:isoamyl alcohol (25:24:1 v/v; centrifugation at 14,000 *g*, 5 min, RT), and chloroform:isoamyl alcohol (24:1 v/w; centrifugation at 14,000 *g*, 5 min, RT). DNA was precipitated using pre-chilled isopropanol (gentle inversions of the tube, centrifugation at 18,000 *g*, 15 min, 4 °C). The resulting pellet was washed with pre-chilled ethanol (70% v/v), before stored in Tris-HCl (100 mM, pH 8) buffer. DNA concentration was determined with NanoDrop (Thermo Scientific), and DNA with A230:260:280 ≈ 1.0:2.0:1.0 was considered appropriate for sequencing. Total RNA was isolated using the RNeasy Plant Mini Kit (Qiagen) following directions of the manufacturer. RNA quality and concentration were determined using Agilent 2100 BioAnalyzer.

### Genome sequence data generation and *de novo* genome assembly

All genome sequence data generated for the five *Symbiodinium* isolates are detailed in Supplementary Table 7. Short-read sequence data (2 × 150 bp reads, insert length 350 bp) were generated using paired-end libraries on the Illumina HiSeq 2500 and 4000 platforms at the Australian Genome Research Facility (Melbourne) and the Translational Research Institute Australia (Brisbane). For all samples, except for *S. pilosum* CCMP2461, an additional paired-end library (insert length 250 bp) was designed such that the read-pairs of 2 × 150 bp would overlap. Quality assessment of the raw paired-end data was done with FastQC v0.11.5, and subsequent processing with Timmomatic v0.36^40^. To ensure high-quality read data for downstream analyses, the paired-end mode of Trimmomatic was run with the settings: ILLUMINACLIP:[AdapterFile]:2:30:10 LEADING:30 TRAILING:30 SLIDINGWINDOW:4:25 MINLEN:100 AVGQUAL:30; CROP and HEADCROP were run (prior to LEADING and TRAILING) when required to remove read ends with nucleotide biases. Genome size and sequence read coverage were estimated from the trimmed read pairs based on *k*-mer frequency analysis (Supplementary Table 2) as counted with Jellyfish v2.2.6; proportion of the single-copy regions of the genome and heterozygosity were computed with GenomeScope v1.0^41^. *De novo* genome assembly was performed for all isolates with CLC Genomics Workbench v7.5.1 (qiagenbioinformatics.com) at default parameters, and using the filtered read pairs and single-end reads. The genome assemblies of *S. microadriatricum* 04-503SCI.03, *S. microadriaticum* CassKB8, *S. linucheae* CCMP2456 and *S. pilosum* CCMP2461 were further scaffolded with transcriptome data (see below) using L_RNA_scaffolder^42^. Short sequences (<1000 kbp) were removed from the assemblies.

### Removal of putative microbial contaminants

To identify putative sequences from bacteria, archaea and viruses in the genome scaffolds, we followed the approach of Chen *et al.*^15^. In brief, we first searched the scaffolds (BLASTn) against a database of bacterial, archaeal and viral genomes from RefSeq (release 88), and identified those with significant hits (*E* ≤ 10^−20^ and bit score ≥ 1000). We then examined the sequence cover of these regions in each scaffold, and identified the percentage (in length) contributed by these regions relative to the scaffold length. We assessed the added length of implicated genome scaffolds across different thresholds of percentage sequence cover in the alignment, and the corresponding gene models in these scaffolds as predicted from available transcripts (see below) using PASA v2.3.3^43^, with a modified script (github.com/chancx/dinoflag-alt-splice) that recognises an additional donor splice site (GA), and TransDecoder v5.2.0^43^. Any scaffolds with significant bacterial, archaeal or viral hits covering ≥5% of its length was considered as a putative contaminant and removed from the assembly (Supplementary Figure 10). Additionally, the length of the remaining scaffolds was plotted against their G+C content; scaffolds (>100 kbp) with irregular G+C content (in this case, G+C ≤45% or ≥60%) were considered as putative contaminant sequences and removed (Supplementary Figure 11).

### Generation and assembly of transcriptome data

We generated transcriptome sequence data for the S*ymbiodinium* isolates, except for *S. necroappetens* CCMP2469 for which the extraction of total RNAs failed (Supplementary Table 8). Short-read sequence data (2 × 150 bp reads) were generated using paired-end libraries on the Illumina NovaSeq 6000 platform at the Australian Genome Research Facility (Melbourne). Quality assessment of the raw paired-end data was done with FastQC v0.11.4, and subsequent processing with Trimmomatic v0.35^40^. To ensure high-quality read data for downstream analyses, the paired-end mode of Trimmomatic was run with the settings: HEADCROP:10 ILLUMINACLIP:[AdapterFile]:2:30:10 CROP:125 SLIDINGWINDOW:4:13 MINLEN:50. The surviving read pairs were further trimmed with QUADTrim v2.0.2 (bitbucket.org/arobinson/quadtrim) with the flags -*m2* and -*g* to remove homopolymeric guanine repeats at the end of the reads (a systematic error of Illumina NovaSeq 6000).

Transcriptome assembly was performed with Trinity v2.1.1^44^ in two modes: *de novo* and genome-guided. *De novo* transcriptome assembly was done using default parameters and the trimmed read pairs. For genome-guided assembly, high-quality read pairs were aligned to their corresponding *de novo* genome assembly (prior to scaffolding) using Bowtie 2 v2.2.7^45^. Transcriptomes were then assembled with Trinity in the genome-guided mode using the alignment information, and setting the maximum intron size to 100,000 bp. Both *de novo* and genome-guided transcriptome assemblies from each of the four samples were used for scaffolding (see above) and gene prediction (see below) in their corresponding genome.

### Gene prediction and function annotation

We adopted the same comprehensive *ab initio* gene prediction approach reported in Chen *et al.*^15^, using available genes and transcriptomes of Symbiodiniaceae as supporting evidence. A *de novo* repeat library was first derived for the genome assembly using RepeatModeler v1.0.11 (repeatmasker.org/RepeatModeler). All repeats (including known repeats in RepeatMasker database release 20180625) were masked using RepeatMasker v4.0.7 (repeatmasker.org).

As direct transcript evidence, we used the *de novo* and genome-guided transcriptome assemblies from Illumina short-read sequence data (see above). For *S. necroappetens* CCMP2469, we used transcriptome data of the other four *Symbiodinium* isolates for gene prediction, as well as other available transcriptome datasets of *Symbiodinium*: *S. microadriaticum* CassKB8^46^, *S. microadriaticum* CCMP2467^10^, *S. tridacnidorum* Sh18^12^, and *S. tridacnidorum* CCMP2592 and *S. natans* CCMP2548^8^. We also combined the *S. microadriaticum* CassKB8 transcriptome data generated here with those from a previous study^46^. We concatenated all the transcript datasets per sample, and vector sequences were discarded using SeqClean (sourceforge.net/projects/seqclean) based on shared similarity to sequences in the UniVec database build 10.0. We used PASA v2.3.3^43^, customised to recognise dinoflagellates alternative splice donor sites (github.com/chancx/dinoflag-alt-splice), and TransDecoder v5.2.0^43^ to predict CDS. These CDS were searched (BLASTp, *E* ≤ 10^−20^) against a protein database that consists of RefSeq proteins (release 88) and a collection of available and predicted proteins (using TransDecoder v5.2.0^43^) of Symbiodiniaceae (total of 111,591,828 sequences; Supplementary Table 9). We used the *analyze_blastPlus_topHit_coverage.pl* script from Trinity v2.6.6^44^ to retrieve only those CDS having an alignment >70% to a protein (*i.e.* nearly full-length) in the database for subsequent analyses.

The near full-length gene models were checked for transposable elements (TEs) using HHblits v2.0.16 (probability = 80% and *E*-value = 10^−5^), searching against the JAMg transposon database (sourceforge.net/projects/jamg/files/databases), and TransposonPSI (transposonpsi.sourceforge.net). Gene models containing TEs were removed from the gene set, and redundancy reduction was conducted using cd-hit v4.6^47,48^ (ID = 75%). The remaining gene models were processed using the *prepare_golden_genes_for_predictors.pl* script from the JAMg pipeline (altered to recognise GA donor splice sites; jamg.sourceforge.net). This script produces a set of “golden genes” that were used as training set for the *ab initio* gene-prediction tools AUGUSTUS v3.3.1^49^ (customised to recognise the non-canonical splice sites of dinoflagellates; github.com/chancx/dinoflag-alt-splice) and SNAP v2006-07-28^50^. Independently, the soft-masked genome sequences were used for gene prediction using GeneMark-ES v4.32^51^. Swiss-Prot proteins (downloaded on 27 June 2018) and the predicted proteins of Symbiodiniaceae (Supplementary Table 9) were used as supporting evidence for gene prediction using MAKER v2.31.10^52^ protein2genome; the custom repeat library was used by RepeatMasker as part of MAKER prediction. A primary set of predicted genes was produced using EvidenceModeler v1.1.1^53^, modified to recognise GA donor splice sites. This package combined the gene predictions from PASA, SNAP, AUGUSTUS, GeneMark-ES and MAKER protein2genome into a single set of evidence-based predictions. The weightings used for the package were: PASA 10, Maker protein 8, AUGUSTUS 6, SNAP 2 and GeneMark-ES 2. Only gene models with transcript evidence (*i.e.* predicted by PASA) or supported by at least two *ab initio* prediction programs were kept. We assessed completeness by querying the predicted protein sequences in a BLASTp similarity search (*E* ≤ 10^−5^, ≥50% query/target sequence cover) against the 458 core eukaryotic genes from CEGMA^54^. Transcript data support for the predicted genes was determined by BLASTn (*E* ≤ 10^−5^), querying the transcript sequences against the predicted CDS from each genome. Genes for which the transcripts aligned to their CDS with at least 50% of sequence cover and 90% identity were considered as supported by transcript data.

Functional annotation of the predicted genes was conducted based on sequence similarity searches against known proteins following the same approach as Liu *et al.*^11^, in which the predicted protein sequences were first searched (BLASTp, *E* ≤ 10^−5^, minimum query or target cover of 50%) against the manually curated Swiss-Prot database, and those with no Swiss-Prot hits were subsequently searched against TrEMBL (both databases from UniProt, downloaded on 27 June 2018). The best UniProt hit with associated GO terms (geneontology.org) was used to annotate the query protein with those GO terms using the UniProt-GOA mapping (downloaded on 03 June 2019). Pfam domains^55^ were searched in the predicted proteins of all samples using PfamScan^56^ (*E* ≤ 0.001) and the Pfam-A database (release 30 August 2018)^55^.

### Comparison of genome sequences and analysis of conserved synteny

We compared the genome data of 15 isolates in Order Suessiales (Supplementary Table 1): the five for which we generated genome assemblies in this study (*S. microadriaticum* CassKB8, *S. microadriaticum* 04-503SCI.3, *S. necroappetens* CCMP2469, *S. linucheae* CCMP2456 and *S. pilosum* CCMP2461), three generated by Shoguchi and collaborators (*B. minutum*, *S. tridacnidorum* Sh18 and *Cladocopium* sp. C92)^12,13^, two from González-Pech *et al.* (*S. tridacnidorum* CCMP2592 and *S. natans* CCMP2548)^8^, two from Liu *et al.* (*C. goreaui* and *F. kawagutii*)^11^, two from Stephens *et al.* (*P. glacialis* CCMP1383 and CCMP2088)^14^, and one from Aranda *et al.* (*S. microadriaticum* CCMP2467^10^. Genes were consistently predicted from all genomes using the same workflow^8,14,15^.

Whole-genome sequence alignment was carried out for all possible genome pairs (225 combinations counting each genome as both reference and query) with nucmer v4.0.0^57^, using anchor matches that are unique in the sequences from both reference and query sequences (--*mum*). Here, the similarity between two genomes was assessed based on the proportion of the total bases in the genome sequences of the query that aligned to the reference genome sequences (*Q*) and the average percent identity of one-to-one alignments (*i.e.* the reciprocal best one-to-one aligned sequences for the implicated region between the query and the reference; *I*). For example, if two genomes are identical, both *Q* and *I* would have a value of 100%. Filtered read pairs (see above, Supplementary Table 7) from all isolates were aligned to each other’s (and against their own) assembled genome scaffolds using BWA v0.7.13^58^; mapping rates relative to base quality scores were calculated with SAMStat v1.5.1^59^. For each possible genome-pair, we further assessed sequence similarity of the repeat-masked genome assemblies based on the similarity between their *k*-mers profiles. To determine the appropriate *k*-mer size to use, we extracted and counted *k*-mers using Jellyfish v2.2.6^60^ at multiple *k* values (between 11 and 101, step size = 2); *k* = 21 was found to capture an adequate level of uniqueness among these genomes as inferred based on the proportion of distinct and unique *k*-mers^61^ (Supplementary Figure 12). We then computed pairwise *D*_*2*_^*S*^ distances (*d*) for the 15 isolates following Bernard *et al.*^62^. The calculated distances were used to build a NJ tree with Neighbor (PHYLIP v3.697)^63^ at default settings. For deriving an alignment-free similarity network, pairwise similarity was calculated as 10 − *d*^64^.

To assess conserved synteny, we identified collinear syntenic gene blocks common to each genome pair based on the predicted genes and their associated genomic positions. Following Liu *et al.*^11^, we define a syntenic gene block as a region conserved in two genomes in which five or more genes are coded in the same order and orientation. First, we concatenated the sequences of all predicted proteins to conduct all-*versus*-all BLASTp (*E* ≤ 10^−5^) searching for similar proteins between each genome pair. The hit pairs were then filtered to include only those where the alignment covered at least half of either the query or the matched protein sequence. Next, we ran MCScanX^65^ in inter-specific mode (−*b 2*) to identify blocks of at least five genes shared by each genome pair. We independently searched for collinear syntenic blocks within each genome (*i.e.* duplicated gene blocks). Likewise, we conducted a BLASTp (*E* ≤ 10^−5^) to search for similar proteins within each genome; the hit pairs were filtered to include only those where the alignment covered at least half of either the query or the matched protein sequence. We then ran MCScanX in intra-specific mode (−*b 1*).

### Genic features, gene families and function enrichment

We examined variation among the predicted genes for all Suessiales isolates with a Principal Component Analysis (PCA; Fig. 3A) using relevant metrics (Supplementary Table 4), following Chen *et al.*^15^. We calculated G+C content in the third position of synonymous codons and effective number of codons used (*Nc*) with CodonW (codonw.sourceforge.net) for complete CDS (defined as those with both start and stop codons) of all isolates. Groups of homologous sequences from all genomes were inferred with OrthoFinder v2.3.1^66^ and considered as gene families. A rooted species tree was inferred using 28,116 families encompassing at least 4 genes from any isolate using STAG^67^ and STRIDE^68^, following the standard OrthoFinder pipeline.

GO enrichment of genes in families core to Symbiodiniaceae and *Symbiodinium* (defined as those common to all isolates in, and exclusive to, each group) was conducted using the topGO Bioconductor package^69^ executed in R v3.5.1, implementing Fisher’s Exact test and the ‘elimination’ method; the GO terms associated to the genes of all isolates surveyed here were used as background to compare against. We considered a *p* ≤ 0.01 as significant.

## Supporting information

Supplementary Figures 1 through 12

Supplementary Tables 1 through 9

## Acknowledgements

R.A.G.P. was supported by an International Postgraduate Research Scholarship and a University of Queensland Centenary Scholarship. This work was supported by two Australian Research Council grants (DP150101875 awarded to M.A.R., C.X.C. and D.B., and DP190102474 awarded to C.X.C. and D.B.), and the computational resources of the National Computational Infrastructure (NCI) National Facility systems through the NCI Merit Allocation Scheme (Project d85) awarded to C.X.C. and M.A.R. We thank Mary Alice Coffroth for her generosity in providing access to the two *S. microadriaticum* cultures used in this study, and Guillaume Bernard for his advice and assistance in interpreting the alignment-free phylogenetic tree and network.

## Author contributions

R.A.G.P., M.A.R. and C.X.C. conceived the study; R.A.G.P., Y.C., T.G.S., S.S., A.R.M., D.B., M.A.R. and C.X.C. designed the analyses and interpreted the results; C.X.C. maintained the dinoflagellate cultures; C.X.C. and A.R.M. extracted biological materials for sequencing; R.A.G.P., Y.C., T.S., S.S. and R.L. conducted all computational analyses. R.A.G.P. prepared all figures and tables, and prepared the first draft of the manuscript; all authors wrote, reviewed, commented on and approved the final manuscript.

## Data availability

The assembled genomes, predicted gene models and proteins from *S. microadriaticum* CassKB8, *S. microadriaticum* 04-503SCI.03, *S. necroappetens* CCMP2469, *S. linucheae* CCMP245 and *S. pilosum* CCMP2461 are available at cloudstor.aarnet.edu.au/plus/s/095Tqepmq2VBztd.

