## Supplementary Figures 1 through 12 for "Genomes of Symbiodiniaceae reveal extensive sequence divergence but conserved functions at family and genus levels"

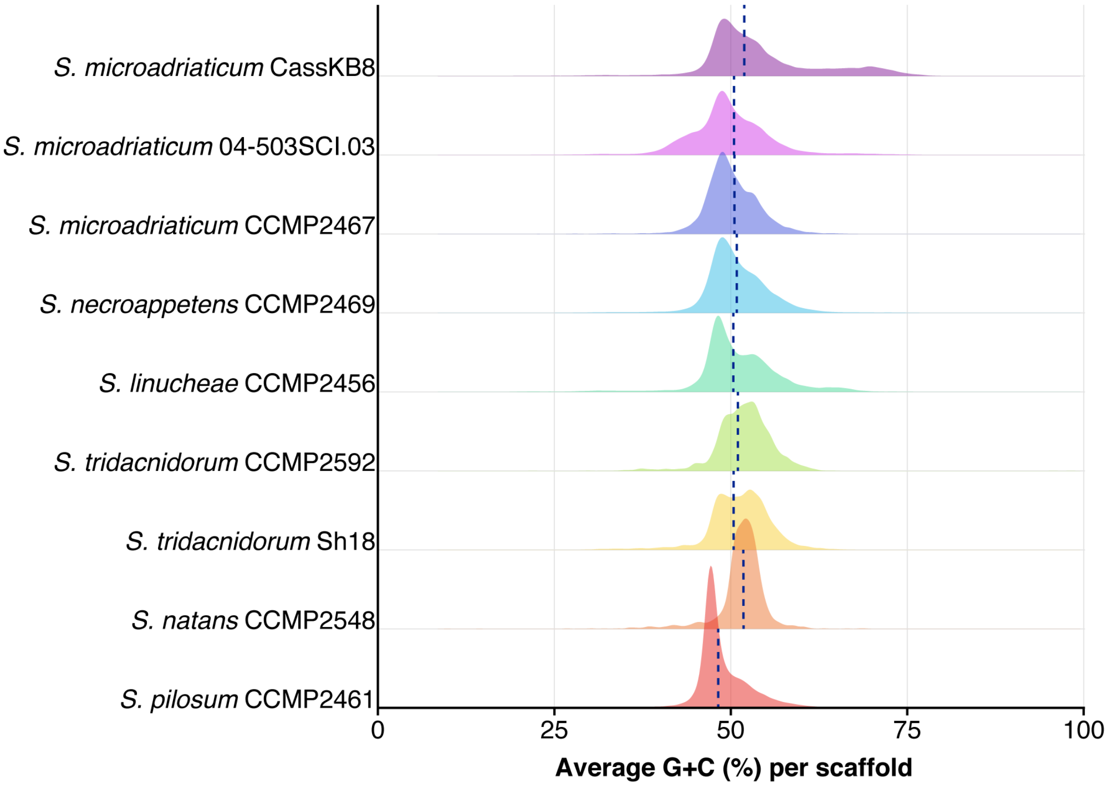


**Supplementary Figure 1** Distribution of per-scaffold G+C content for each assembled genome of *Symbiodinium*.

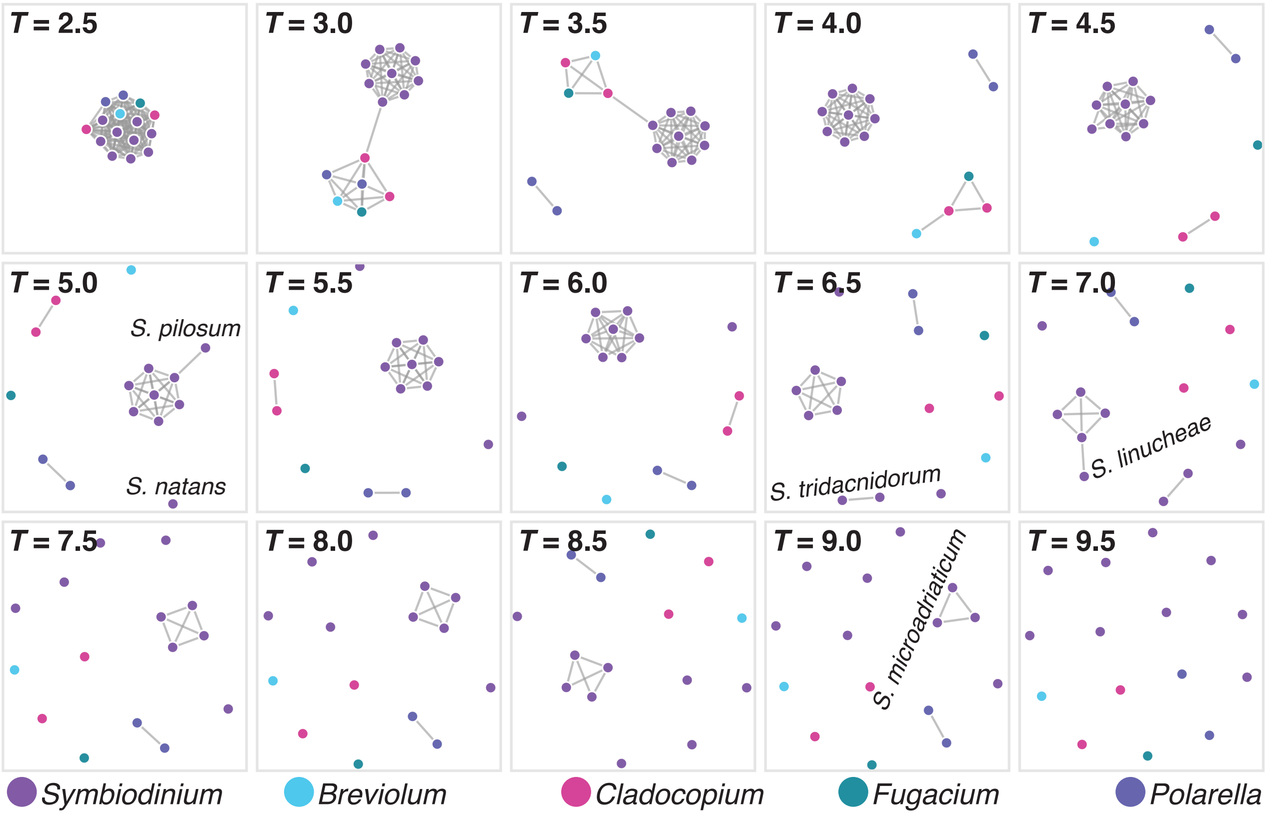


**Supplementary Figure 2** Similarity network constructed based on shared 21-mers among the genome sequences of the Suessiales isolates. Each panel shows the network at a similarity threshold (*T*), at edges (connexions) with a similarity value below the threshold are removed; see Bernard *et al.* (2018) for more detail. Data points representing each dataset are coloured following the bottom legend. *S. microadriaticum* isolates are the last *Symbiodinium* clique at $T = 9.0$.

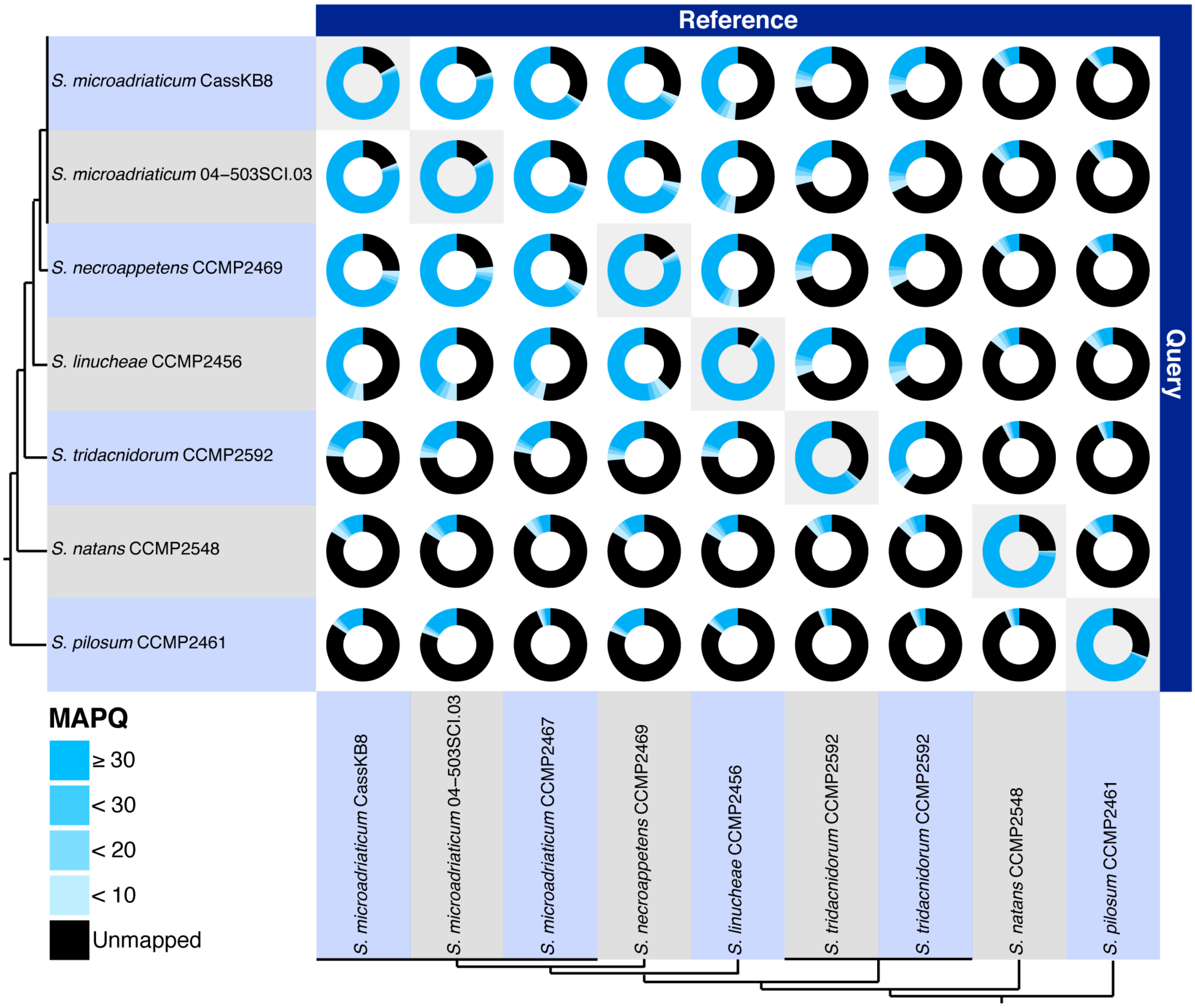


**Supplementary Figure 3** Mapping rate of filtered paired reads that we generated for each *Symbiodinium* isolate against the assembled genomes of itself (grey background) and of all other *Symbiodinium* isolates. The tree topologies on the left and bottom indicate the known phylogenetic relationship (LaJeunesse *et al.*, 2018) among the isolates.

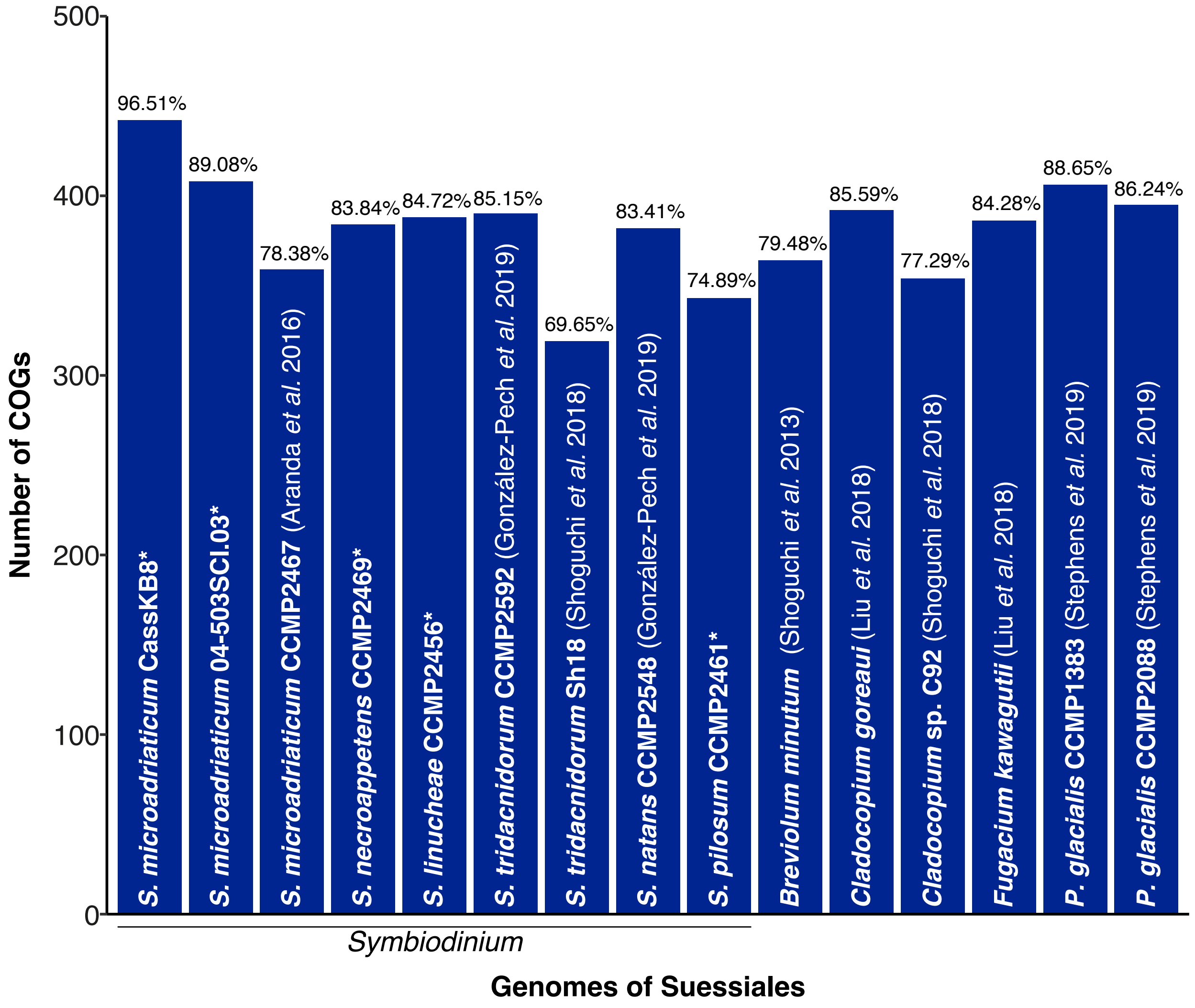


**Supplementary Figure 4** Recovery of 458 conserved eukaryote genes among the predicted genes in each genome, based on CEGMA Clusters of Orthologous Groups. Isolates for which genome data were generated in this study are indicated with an asterisk. The percentage of recovered CEGMA genes is shown for each bar.

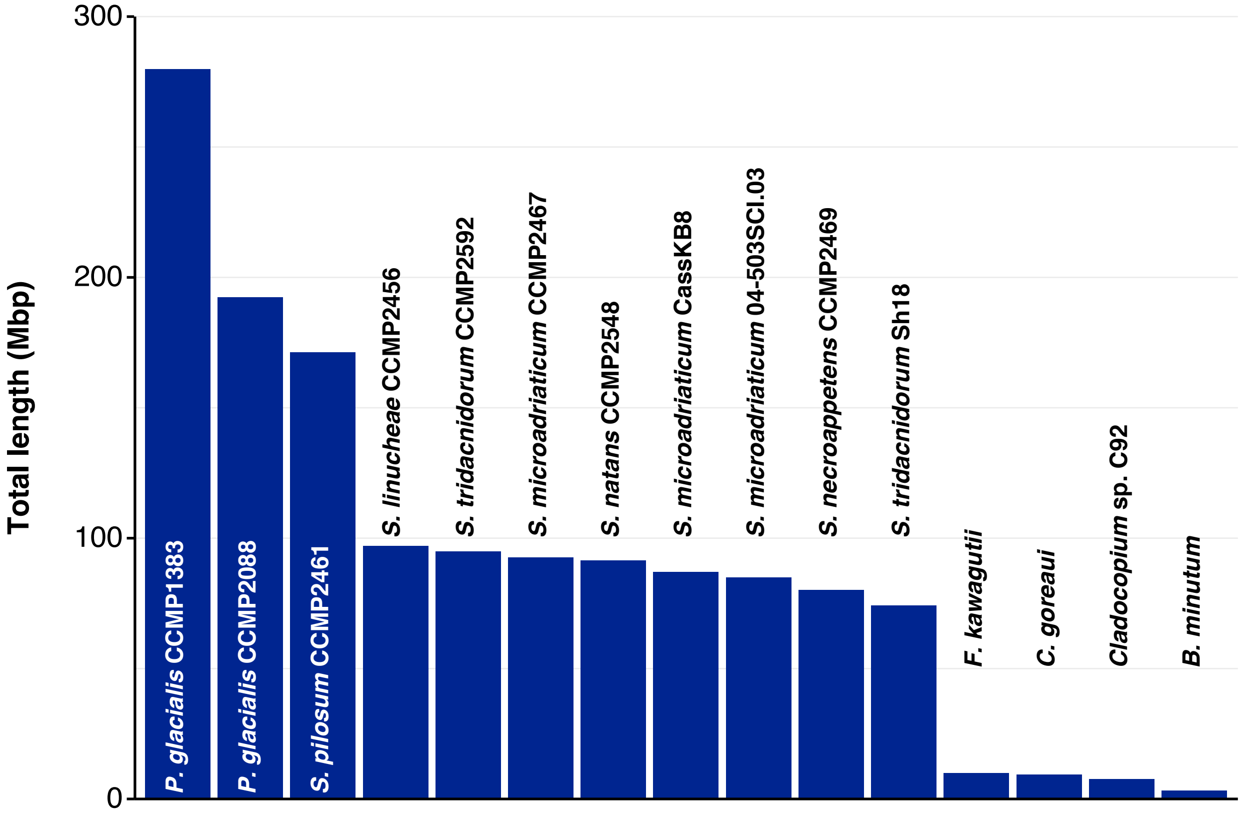


**Supplementary Figure 5** Total length of each analysed genome that comprises LINEs, sorted in decreasing order.

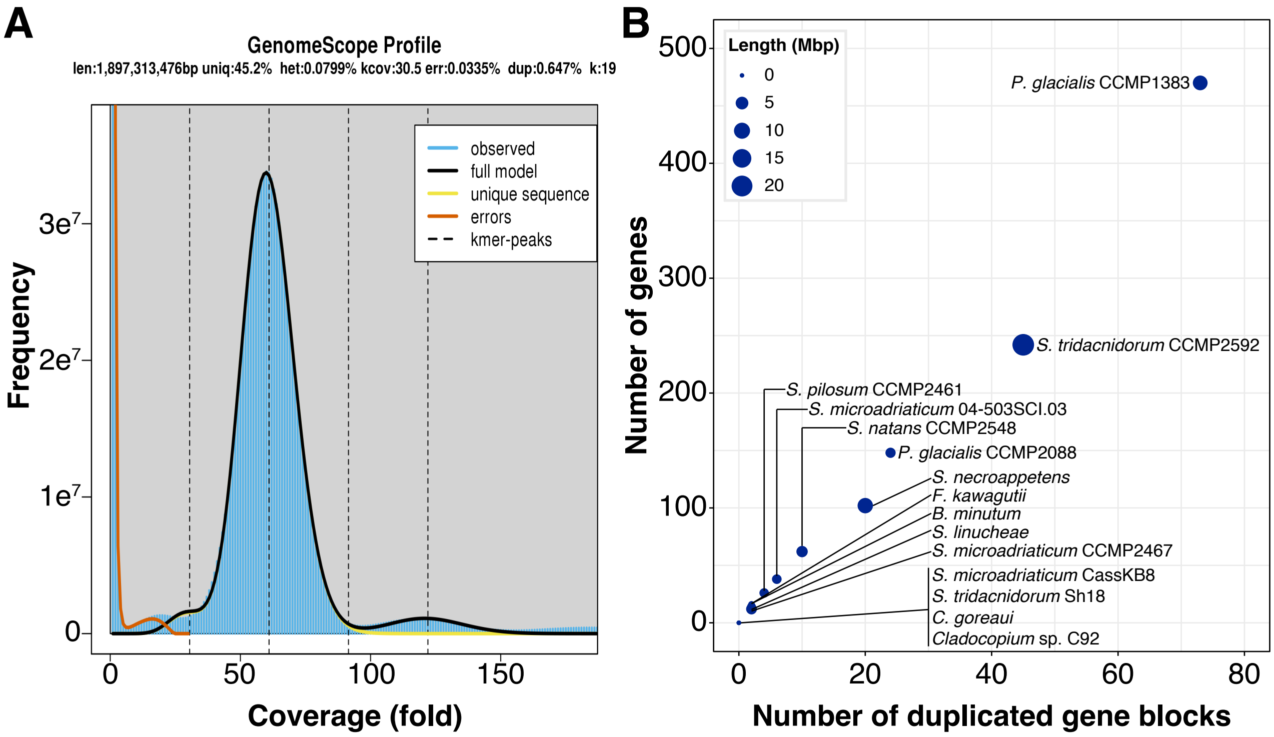


**Supplementary Figure 6 (A)** GenomeScope profile based on 19-mers. **(B)** Number of the duplicated gene blocks found within each genome (*x*-axis) against the number of implicated genes in those blocks (*y*-axis). The size of the dot is proportional to the added sequence length comprising the duplicated gene blocks, as shown in the top-left legend.

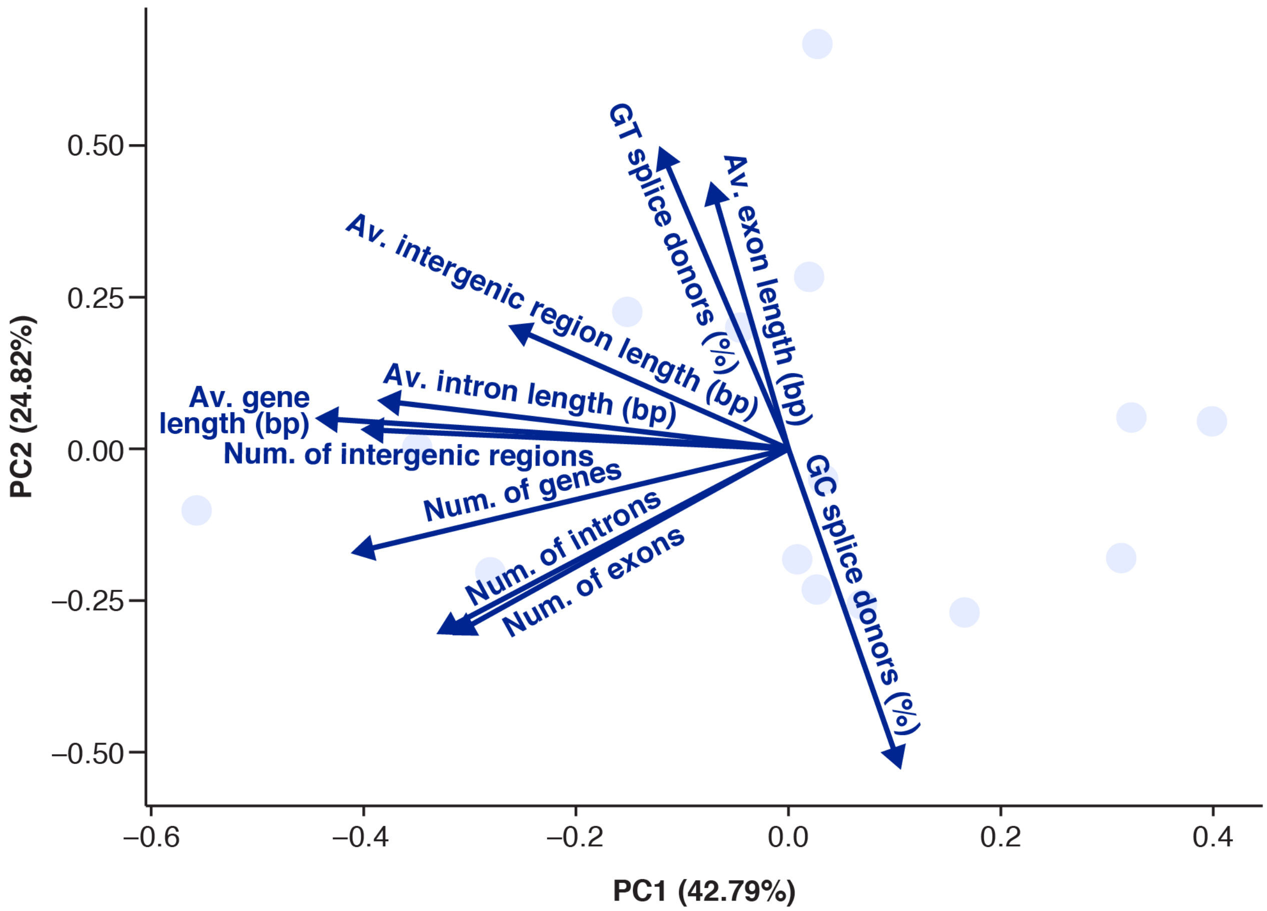


**Supplementary Figure 7** Loading plot showing the contribution of the distinct gene metrics employed for the PCA (Supplementary Table 4, Fig. 3A) to PC1 and to PC2.

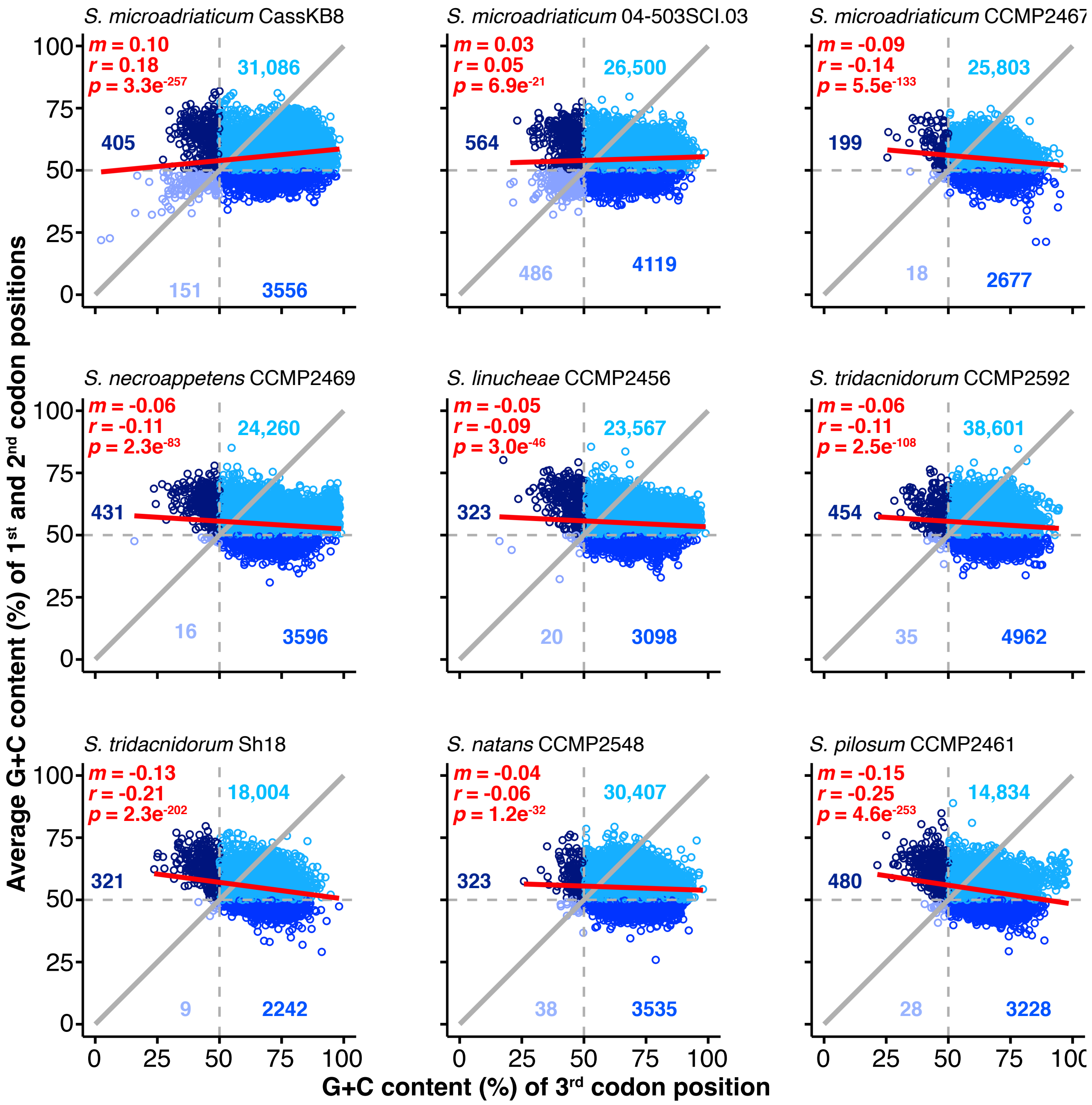


**Supplementary Figure 8** G+C content of third codon position (*x*-axis) against average G+C content of first and second positions (*y*-axis) of full-length CDS. The grey diagonal line indicates the values where the nucleotide composition is the same in both metrics, indicative of neutral evolution. The red line represents the regression line estimated from the CDS data; the corresponding slope (*m*), Pearson's correlation coefficient (*r*) and statistical significance (*p*) are shown in red. To highlight overall patterns of nucleotide composition, the plot is split into four quadrants with coordinates at 50% from both axes. Dots falling into each quadrant are coloured in a different shade of blue; the corresponding number of CDS for each quadrant is shown.

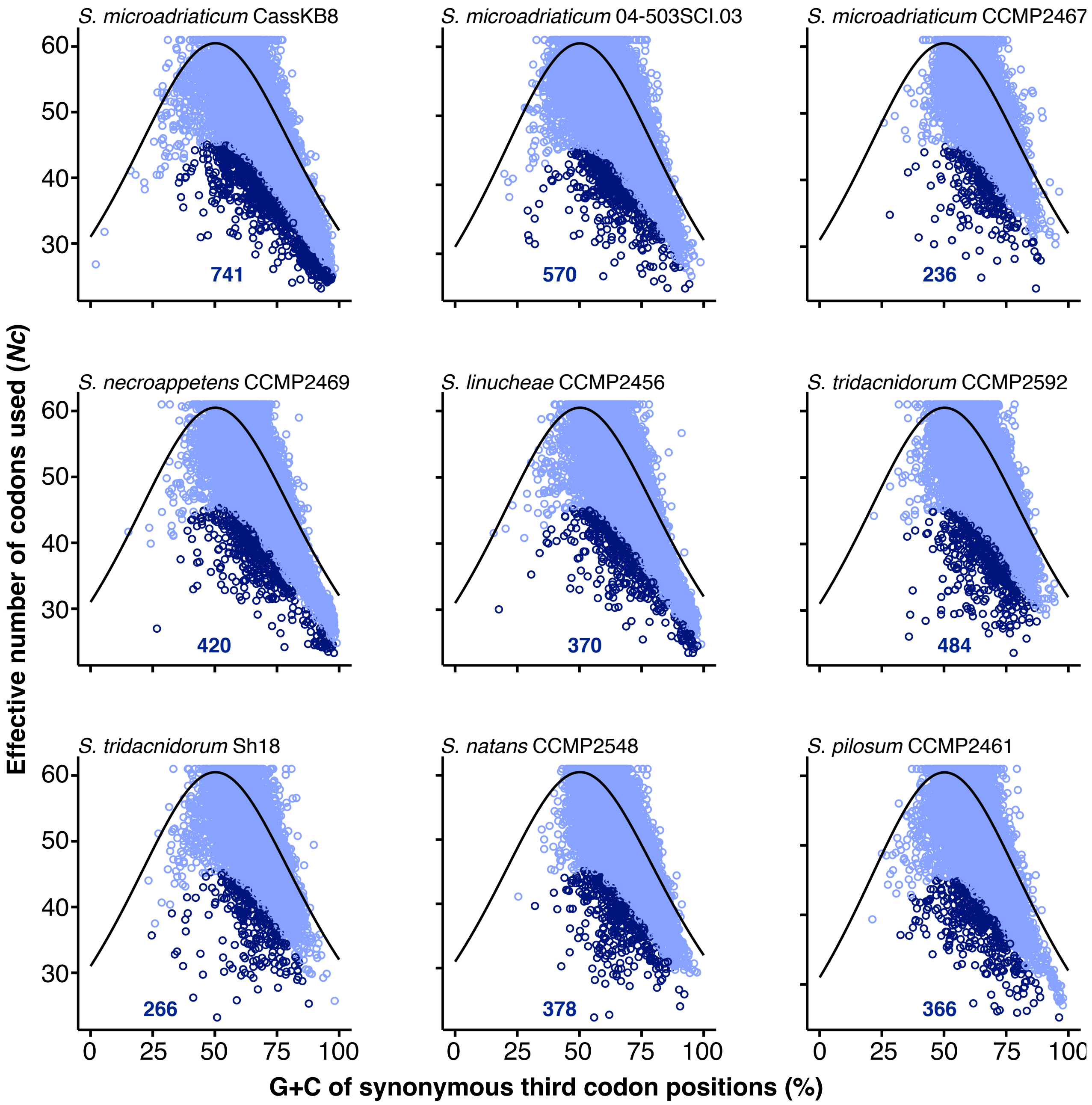


**Supplementary Figure 9** Effective number of codons used (*Nc*, *y*-axis) as a function of the G+C content of synonymous third codon positions (*x*-axis). The curve line represents the neutral expectation of *Nc*. CDS with a *Nc* 25% smaller than the expected are considered to display strong codon usage preference and are highlighted in a darker shade of blue; the corresponding CDS count is shown in each graph.

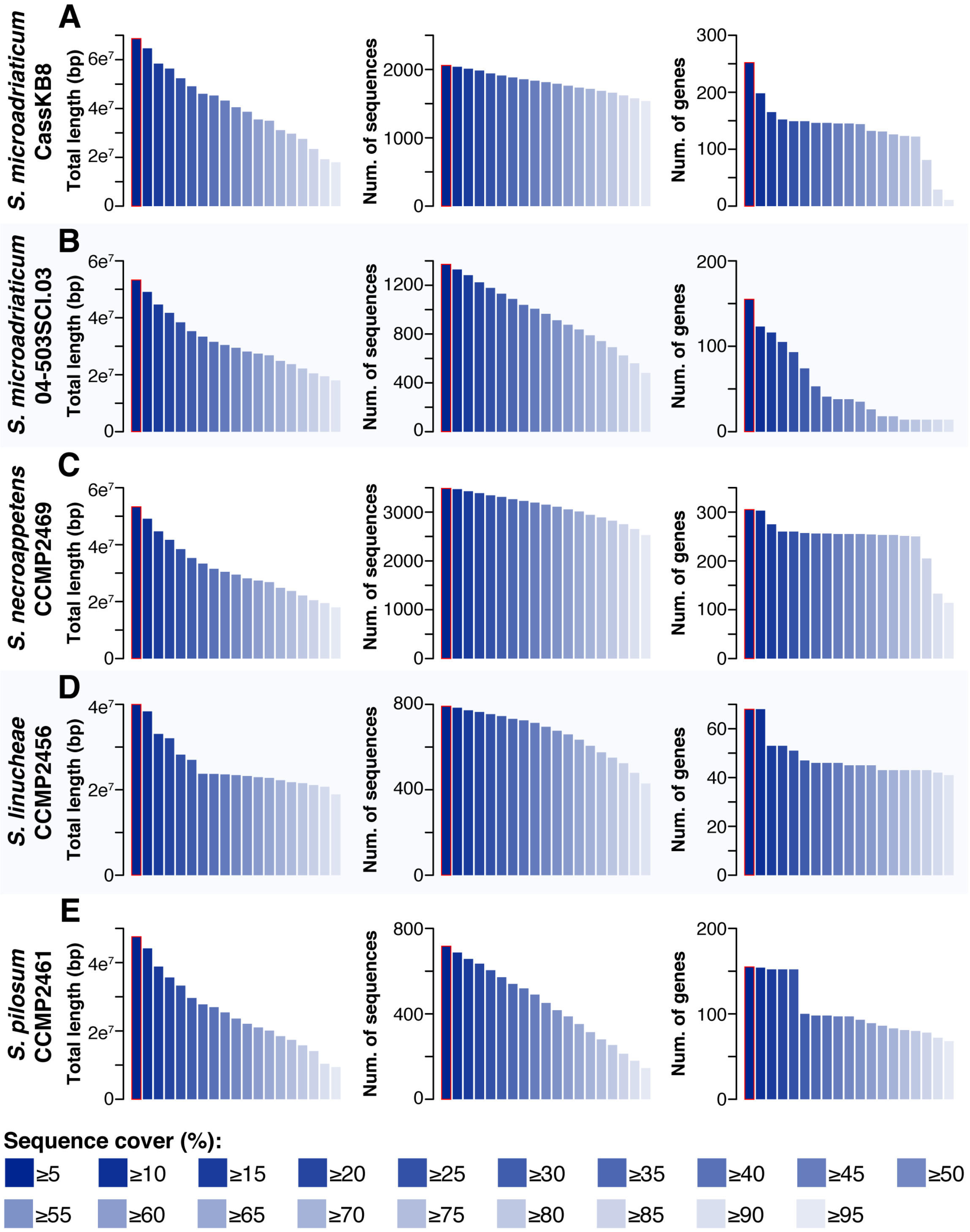


**Supplementary Figure 10** Identification of putative contaminant sequences in each assembled genome of *Symbiodinium*. The total length of the genome sequences with shared significant similarity to known bacterial, archaeal and viral genome sequences (left panels), the number of these implicated sequences (middle panels), and the number of genes located in these implicated sequences (right panels) are shown, across different thresholds of the percentage of sequence cover relative to aligned regions. The threshold of 5% (bars with red outline) is the chosen threshold in this analysis, and the implicated sequences were considered as contaminants and removed from the corresponding assembly.

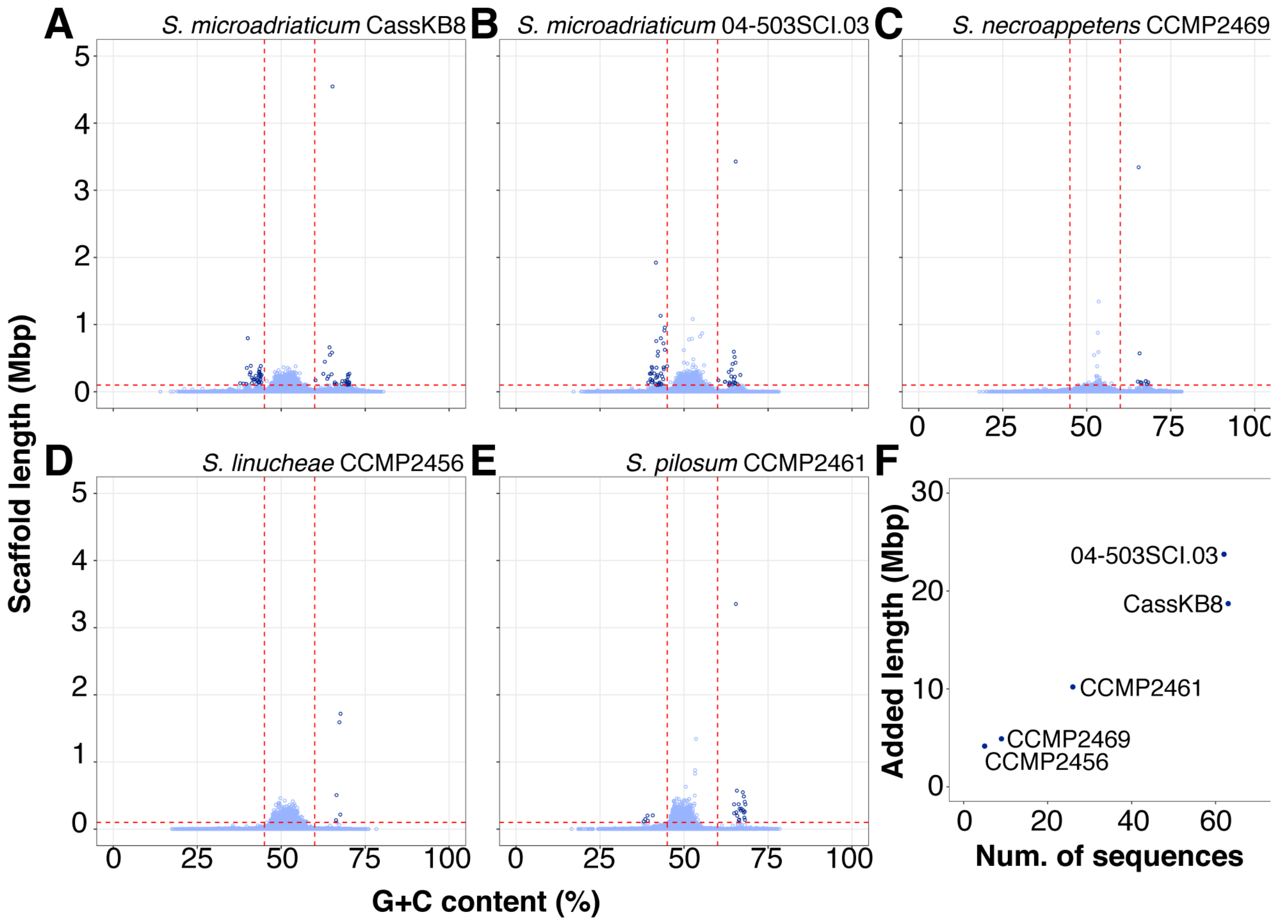


**Supplementary Figure 11 (A-E)** Distribution of G+C content in assembled genome scaffolds relative to the scaffold lengths, shown for each *Symbiodinium* genome generated in this study. Data points in a darker shade of blue represent scaffolds that were identified as outliers and removed from the assembly. **(F)** Summary of the identified outlier scaffolds relative to their total length.

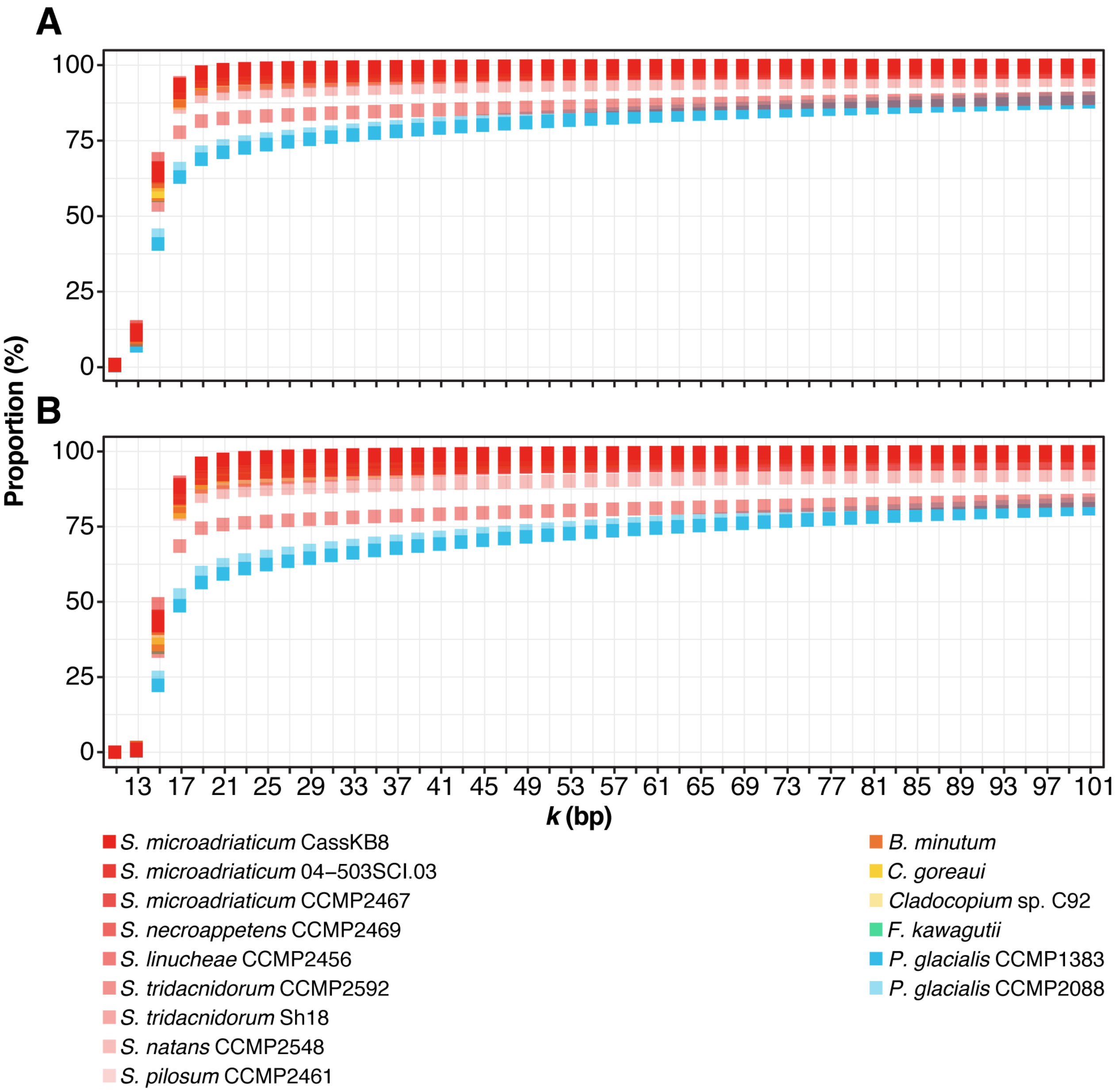


**Supplementary Figure 12** Proportion of distinct **(A)** and unique **(B)** *k*-mers at different *k*-values in each isolate (coloured according to the bottom legend).
